# Developmental senescence profiling reveals Eda2r as a common mediator in the core senescence program

**DOI:** 10.1101/2025.10.30.685553

**Authors:** Annabelle Klein, Daniel Sampaio Gonçalves, Ludivine Dulac, Tania Knauer-Meyer, Matej Durik, Adelyne S.L. Chan, Marie Christine Birling, Muriel Rhinn, Masashi Narita, William M. Keyes

## Abstract

Cellular senescence is a distinct state of stable cell-cycle arrest coupled to a dynamic secretory phenotype. Although transient senescence contributes to embryonic development and regeneration, its relationship to the chronic forms associated with damage and aging is unclear. Here, we generated a p21-reporter mouse to isolate and transcriptionally profile senescent cells in Apical Ectodermal Ridge of the developing limb. Comparative analysis of these cells with diverse *in vitro* and *in vivo* senescence models revealed a conserved gene program encompassing canonical markers and regulators, with previously unassociated factors. Among these, ectodysplasin A2 receptor (Eda2r) emerged as a critical modulator required for senescence-associated transcriptional and functional features. Our findings define a shared molecular framework linking developmental and pathological senescence and identify conserved effectors such as Eda2r as potential targets for therapeutic modulation of senescent cell function.

## INTRODUCTION

Cellular senescence is a state of stable cell cycle arrest associated with the upregulation of cell cycle inhibitors, a robust secretory phenotype known as the SASP (senescence-associated secretory phenotype), and, in some contexts, features of macromolecular damage^1–3^. In aged or damaged tissues, senescent cells accumulate and contribute to tissue dysfunction, chronic inflammation, and age-related disease ^4,5^. Functional studies using genetic or pharmacological approaches to eliminate or disrupt senescent cells have demonstrated a contributory role in aging and diverse pathologies, including neurodegeneration, fibrosis, and cancer^1^. Senescence-targeting strategies have typically relied on mouse models engineered to eliminate cells expressing canonical markers such as p16 (*Cdkn2a*) or p21 (*Cdkn1a*)^6–8^. Additional approaches have involved senolytic compounds that induce apoptosis in senescent cells, or senomorphic agents that suppress the SASP^9,10^. More recently, antibody-based strategies targeting senescence-associated surface proteins have been developed to block signaling or deliver cytotoxic payloads^11–13^. Collectively, these interventions support the idea that selective disruption of senescent cells or their secretory output can confer therapeutic benefit in aging and disease contexts.

However, senescence also plays important physiological roles in embryonic development, tissue regeneration, and wound healing^8,14–17^. In these contexts, the transient appearance of senescent cells and the SASP promotes tissue remodeling, cell plasticity, and repair, before being efficiently cleared by the immune system^2,18^. These observations support the view that senescence is a beneficial and tightly regulated process in normal development and regeneration, but becomes dysregulated and persistent in aging and disease.

We previously identified transient populations of senescent cells in the developing embryo, notably in structures such as the apical ectodermal ridge (AER) of the limb and the hindbrain roof plate^15^. These embryonic senescent cells exhibited hallmark features of senescence, including SA-β-gal activity, p21 expression, and a secretory phenotype consistent with the SASP. However, they lacked several features commonly associated with senescent cells in aged or diseased tissues, such as p16 expression, pro-inflammatory cytokines, and markers of DNA-damage. These observations raised the possibility that developmental senescence may represent a more limited or primordial version of the senescence program. Nevertheless, the degree to which developmental senescence resembles or diverges from senescence in pathological or stress-induced contexts has remained unresolved.

Given the shared features observed between developmental and pathological senescence, we hypothesized that common molecular programs shared between these states may exist. To test this, we sought to directly compare developmental senescence with various established senescence models, with the aim of identifying conserved gene signatures and potential regulators. To enable this, we generated a p21-reporter mouse model, allowing for the isolation and transcriptional profiling of senescent cells in the embryo. Focusing on the AER, we isolated p21-expressing cells from E11.5 limbs for RNA sequencing and compared their senescence signature to 18 published datasets. This revealed a conserved set of genes and senotherapeutic targets across models, including known markers such as p21, Cd36, and Dpp4, as well as previously uncharacterized genes consistently upregulated in senescence. Among these, we identified Eda2r as a novel regulator essential for multiple senescence features. These findings define a core developmental senescence signature conserved across contexts and highlight new regulators such as Eda2r for possible therapeutic modulation.

## RESULTS

### Characterization of a new p21 reporter mouse model

To investigate the role of p21 and senescence *in vivo*, we generated a p21-mCherry-CreERT2 reporter mouse model (Fig. 1a and Methods). We could verify the presence of the p21-mCherry-CreERT2 allele by genotyping, and the Mendelian ratio was respected in heterozygous intercrosses for WT (*p21+/+*), heterozygous-reporter (*p21+/m*) and homozygous reporter (*p21m/m*) new-born pups (Extended Data Fig. 1a).

**Figure 1.**
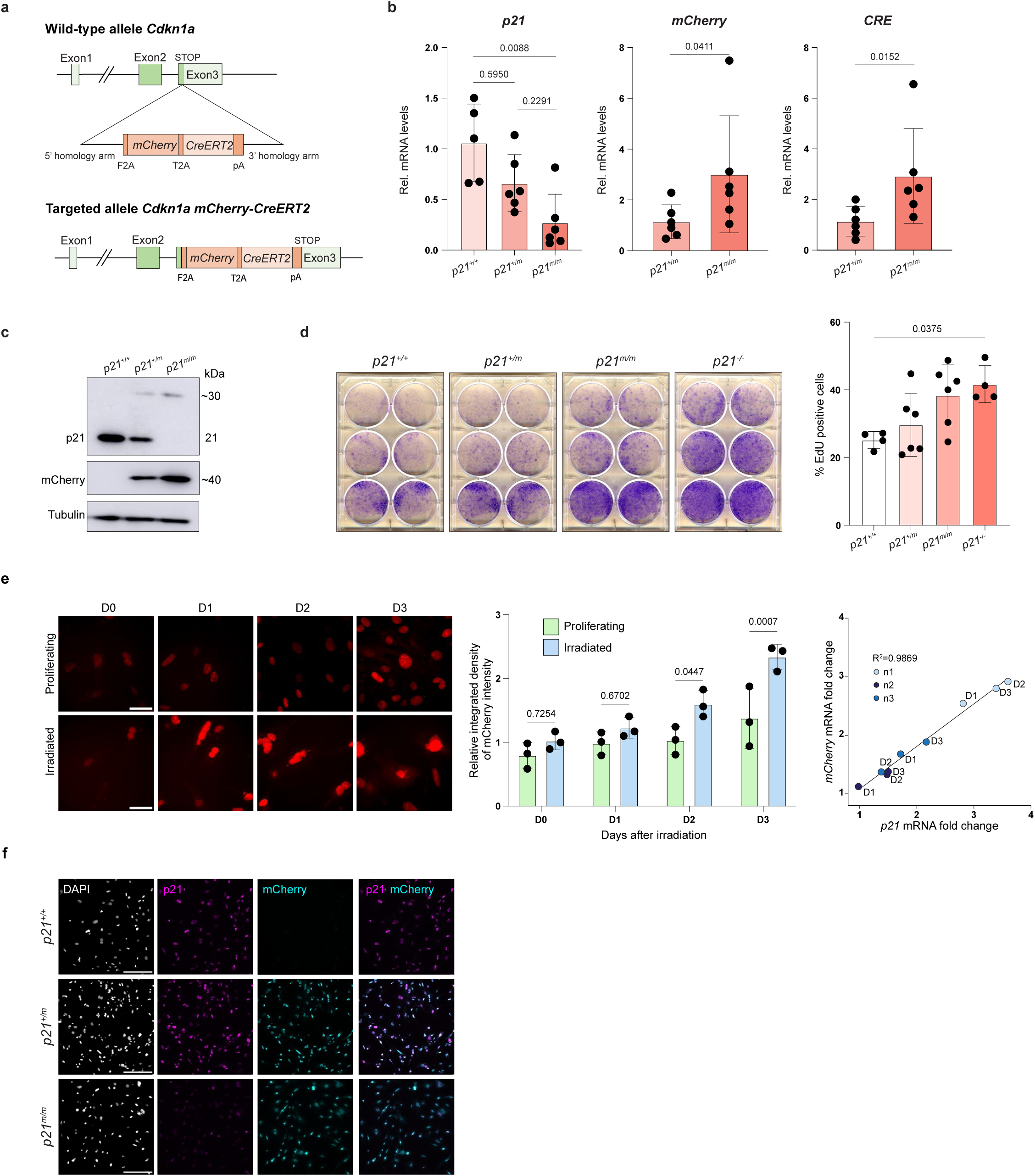
Generation and validation of a new p21-mCherry-CreERT2 reporter model. **a**, Schematic of the insertion strategy of mCherry-CreERT2 cassette within endogenous *Cdkn1a* (*p21*) locus and the obtained genetically modified allele. **b**, Relative mRNA levels of *p21* in proliferating *p21^+/+^*, *p21^+/m^* and *p21^m/m^* mouse dermal fibroblasts (MDFs), and relative mRNA levels of *mCherry* and *Cre* in proliferating *p21^+/m^* and *p21^m/m^*MDFs, normalized to Rplp0 and compared to *p21^+/+^* or *p21^+/m^*respectively. *mCherry* and *Cre* mRNA undetectable in *p21^+/+^*MDFs. n=5-6, Kruskall-Wallis test followed by Dunn’s test for p21, Mann-Whitney tests for mCherry and Cre. **c**, Immunoblotting for p21, mCherry and Tubulin in *p21^+/+^*, *p21^+/m^* and *p21^m/m^* proliferating MDFs, **d**, Left-panel, Focus formation assay showing cell density after 10 days in culture of *p21^+/+^*, *p21^+/m^*, *p21^m/m^* and *p21^-/-^*cells. MDFs were seeded at 2,500 (top row), 5,000 (middle row) and 10,000 (bottom row) cells/wells. Figure is representative of n=4 experiments done on 4 different biological replicates. Right panel, Percentage of EdU positive cells after 2H15 incorporation, in *p21^+/+^*, *p21^+/m^*, *p21^m/m^* and *p21^-/-^* MDFs. n=4-6; one-way ANOVA test followed by Tukey’s multiple comparisons test. **e**, Left, Representative images of mCherry signal in living *p21^+/m^* MDFs, proliferating or following irradiation (D0 to D3 after irradiation); scale bar: 50 µm. Middle, Mean integrated density of mCherry intensity in *p21^+/m^*MDFs proliferating or after irradiation, normalized to a separate control before irradiation, mean of integrated density of 20-300 cells per experiment and per condition, two-way ANOVA test followed by Šídák’s multiple comparisons test. Right, Correlation between mCherry mRNA fold change and p21 mRNA fold change at D1, D2, D3 after irradiation, normalized to D0, n=3. **f**, Immunofluorescence staining for p21 and mCherry in *p21^+/+^, p21^+/m^* and *p21^m/m^* MDFs 3 days after irradiation. Scale bars: 200 µm; n=2-3.

The expression of *p21*, *mCherry* and *Cre* mRNA were first assessed by RT-qPCR in proliferating mouse dermal fibroblasts (MDFs) for each genetic condition (Fig. 1b). Surprisingly, *p21* mRNA levels were approximatively 50% lower in *p21+/m* cells compared to *p21+/+* counterparts, and expression was further decreased in *p21m/m* proliferating MDFs, suggesting that the inserted cassette disrupts p21 gene expression. As expected, *mCherry* and *Cre* mRNA expression were not detectable in *p21+/+* cells but were detectable in increasing levels in *p21+/m* and *p21m/m* cells. Immunoblotting for p21 and mCherry, and immunostaining for p21 in proliferating MDFs confirmed these results (Fig. 1c, Extended Data Fig. 1b). These results highlight that the inserted construct accurately allows expression of mCherry and Cre but disrupts p21 expression at the transcript and protein level.

We then tested whether this disruption is sufficient to affect p21 functions in cells. As p21 is an inhibitor of cell-cycle progression, its decrease or malfunction should increase proliferation. To assess this, we performed a focus formation assay (FFA) on *p21+/+*, *p21* germline knockout (*p21-/-*), *p21+/m* and *p21m/m* MDFs (Fig. 1d). In *p21+/m* cells, density was similar to *p21+/+* cells, but *p21m/m* showed greater density than both these lines, more similar to *p21-/-* MDFs. Similar results were found using an EdU incorporation assay (Fig. 1d). Overall, these results support that p21 expression is disrupted by the genetic construct, but that p*21+/m* cells seem to function like wild-type cells.

We next investigated the functionality of the cassette in an *in vitro* model of induced p21 expression and senescence. For this, MDFs were exposed to 10 Gray (Gy) of X-ray radiation to induce senescence, which was confirmed by increasing p21 mRNA levels in WT cells (Extended Data Fig. 1c). To determine if the expression of mCherry accurately reported levels of p21, we checked the intensity of mCherry in living *p21+/m* MDFs in the same conditions (Fig. 1e). Accordingly, an increase of mCherry signal was observed in living *p21+/m* at D2 and D3 after irradiation, similar to the increase of p21 mRNA levels. Furthermore, the expression of mCherry mRNA was determined to linearly and significantly correlate with the mRNA fold change of p21 in irradiated *p21+/m* cells. These results indicate that mCherry, its signal and mRNA expression, can serve as a good indicator of p21 expression level.

Immunostaining for p21 and mCherry on MDFs 3 days after irradiation showed p21 expression was highly detectable in *p21+/+* and *p21+/m* cells but was dramatically reduced in *p21m/m* cells, and that *p21+/m* and *p21m/m* cells displayed a strong mCherry protein signal, undetectable in WT cells (Fig. 1f). To further determine if p21 function is affected in our model, we checked the expression of *Hmgb2,* a gene repressed by p21, in irradiated MDFs (Extended Data Fig. 1d)^19^. Here, *Hmgb2* mRNA expression was increased in both *p21+/m* and *p21m/m* irradiated MDFs compared to irradiated control. Altogether, these results indicate that the homozygous version of the reporter seems to behave like a p21-hypomorph, including in a p21-high senescent context, and that the wild-type allele of p21 present in heterozygous cells appears sufficient to assure its proper functions. Importantly, this describes the generation and characterization of a new reporter mouse model to study p21 and senescence.

### Isolation and transcriptional profiling of developmental senescent cells

As an aim of this project is to study senescence during embryonic development, we examined expression in living p21+/m embryos. At E10.5, the mCherry pattern perfectly matched the embryonic pattern of p21, as shown by in situ hybridization, notably in the somites, the developing limb and the hindbrain (Fig. 2a). These same structures were also mCherry-positive at E11.5 in p21m/m embryos (Extended Data Fig. 2a). Western blot analysis for p21 and mCherry protein in whole-embryo lysates from E11.5 embryos showed similar results to those found in MDFs, with decreased p21 protein levels in *p21+/m* and absent in *p21m/m*, and with mCherry protein present in both *p21+/m* and *p21m/m* embryos (Extended Data Fig. 2b).

**Figure 2.**
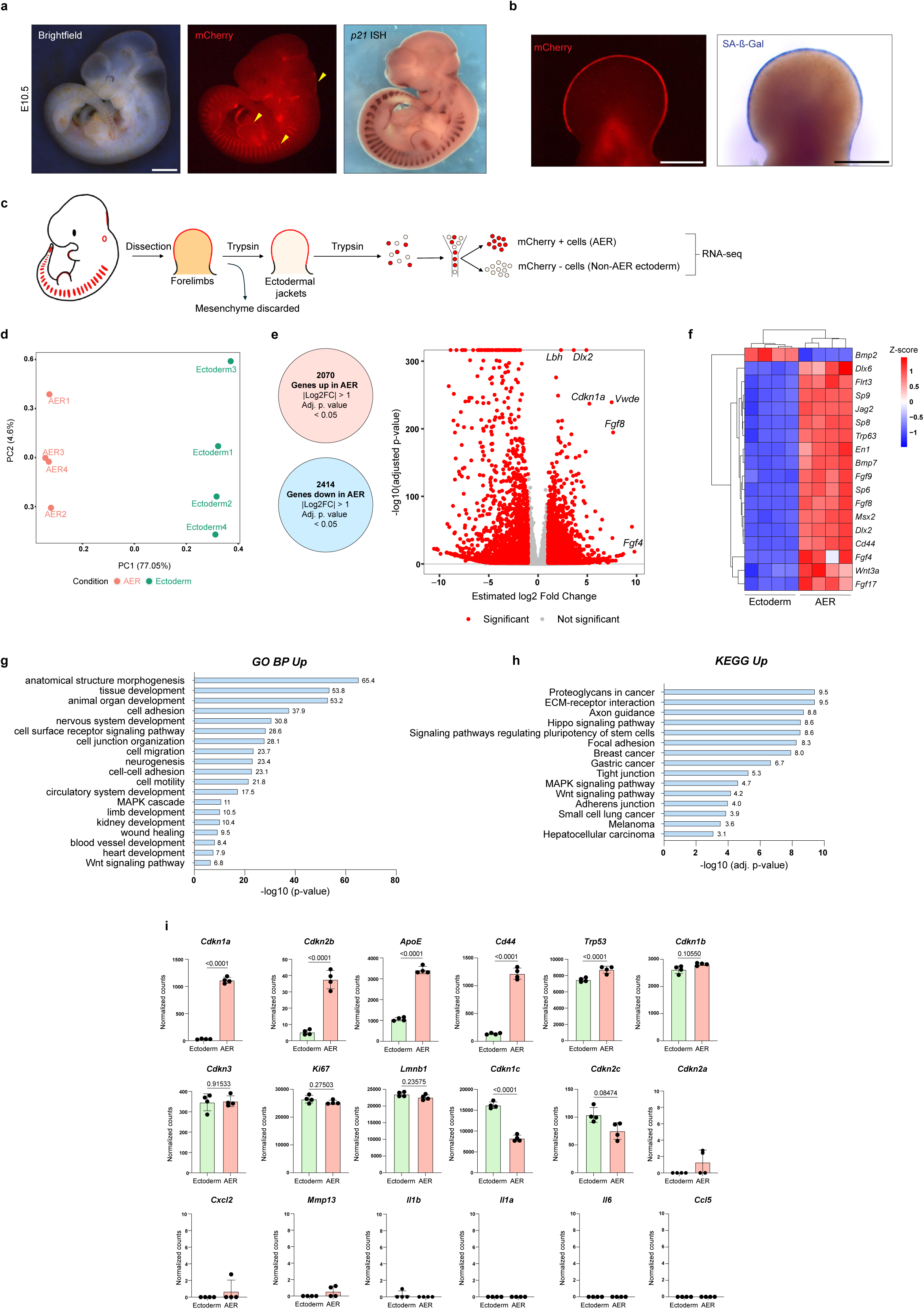
Profiling of developmental senescent cells from the limb apical ectodermal ridge. **a,** Left, pictures of living *p21^+/m^* E10.5 embryo, brightfield and mCherry fluorescence. Arrows denote positive staining in somites, limb and hindbrain. Right, In situ hybridization for p21 at E10.5. Scale bar: 1 mm**. b,** Pictures of live mCherry fluorescence and fixed SA-ß-Gal staining of E11.5 *p21^+/m^*forelimb. Scale bars: 500 µm. **c**, Schematic of the dissection and dissociation protocol used for FACS of mCherry+ and mCherry-forelimb cells, followed by RNA-seq**. d,** Principal component analysis (PCA) of 4 AER and 4 ectoderm samples, from 4 different litters, used for the RNA-seq. **e**, Left, number of differentially expressed genes (DEGs) in the AER, Right, Volcano plot of upregulated and downregulated genes of the AER and ectoderm comparison. Significance determined by absolute log_2_ fold change ≥1 and adjusted p-value <0.05. **f,** Heatmap of normalized reads of selected AER genes in *p21^+/m^* ectoderm and *p21^+/m^* AER. Row scaling. All genes represented here were significantly differentially expressed with an adjusted p-value <0.0001 between ectoderm and AER samples. **g**, Significant selected Gene Ontology (GO) Biological Process (BP) terms of upregulated genes (absolute fold change ≥1.5 and adjusted p-value <0.05) in the AER. **h**, Selected KEGG pathway analysis of upregulated genes (absolute fold change ≥1.5 and adjusted p-value <0.05) in the AER. **i**, Senescent genes normalized reads in ectoderm and AER samples, adjusted p-value not shown for genes whose expression was undetected in all samples of at least one condition.

One goal was to profile senescence in the apical ectodermal ridge (AER) of the developing limb, a well-described signaling center that contributes to the proper development of the limb^20^, which we have previously described as being positive for p21 and senescence markers. In our model, mCherry fluorescence followed previously described SA-ß-Gal patterns, at least for the stages examined, from E10.5 to E12.5 in heterozygous embryos (Extended Data Fig. 2c). And as before, the AER at E11.5 exhibited strong uniform SA-ß-Gal staining, which was matched exactly by the p21-mCherry pattern (Fig. 2b). Importantly, no major phenotypic changes were observed in the embryos. To further ensure that the AER is not disrupted by the presence of the genetic construct, or the decreased levels of p21 in our heterozygous reporter mice, we analyzed the expression of p21 and mCherry and key AER genes, namely *Fgf8*, *Fgf4*, *Wnt3a*, *p63* and *Cd44,* in ectodermal jackets (EJs) of forelimbs from *p21+/+*, *p21+/m* and *p21-/-* embryos (Extended Data Fig. 2d, e, f). As before, p21 levels was decreased in p21+/m compared to WT EJs, albeit not significantly, and mCherry was detected in p21+/m but not in WT EJs (Extended Data Fig. 2e). Importantly, none of the AER genes showed significant disruption between the *p21+/+* and *p21+/m* limbs, while expression was lower in *p21-/-* tissues, consistent with previous findings (Extended Data Fig. 2f). These experiments support that the heterozygous p21-mCherry-CreERT2 reporter mouse is a good model to study p21-positive AER cells.

To molecularly profile the senescent AER cells, we enzymatically dissociated forelimbs from E11.5 *p21+/m* mice, to retrieve the outer ectodermal layer, including the AER. This was then further dissociated into single cell populations, which were sorted to collect the p21-postive AER cells, or p21-negative surface ectoderm, which were then both analyzed by bulk RNA-sequencing (Fig. 2c). PCA analysis showed clear separation of the populations (Fig. 2d), The comparison of *p21+/m* AER with *p21+/m* ectoderm identified a total of 4484 genes that were significantly different, with 2070 being over-expressed in the AER, and 2414 being over-expressed in the ectoderm (Fig. 2e). To further confirm that the *p21+/m* AER samples indeed display an AER gene expression signature, we checked the expression of key genes of this signaling center, including *Fgf8*, *Fgf4*, *Cd44*, *Wnt3a* and others (Fig. 2f), which confirmed all genes were clearly upregulated in the AER samples compared to the ectoderm. Selected pathway analysis for the terms and signatures associated with the differentially expressed genes between the AER and ectoderm showed terms including *cell-adhesion*, *cell-migration*, *neurogenesis*, and signatures related to limb-, kidney-, heart- and blood-development were all associated with the AER genes, consistent with this being a major signaling centre (Fig. 2g, h, Extended Data Fig. 2g, h, i). Interestingly, KEGG analysis linked the AER signature with adhesion (*focal adhesion*, *tight junction*, *adherens junction*), as well as various cancer types (*breast*, *gastric*, *small cell lung*, *melanoma* and *hepatocellular carcinoma*) (Fig. 2h).

Finally, we examined known senescence-associated genes in these samples (Fig. 2i). In agreement with previous microarray profiling^15^, this current dataset highlights key cell-cycle inhibitors are increased in the AER, including *Cdkn1a* (p21), *Cdkn2b* (p15), *Cdkn1b* (p27), while other including *Cdkn1c* (p57) and *Cdkn2c* (p18) were expressed at higher levels in the surface ectoderm. In agreement with previous results, *Cdkn2a* (p16/p19) was either not detectable or expressed at very low reads. Other markers such as *Ki67* and *Lmnb1* were unchanged between samples, and others such as *ApoE* and *Cd44* were expressed more strongly and significantly in the AER. With regards to the SASP and secreted factors produced by this structure, this is discussed in detail in another study ^21^. However, our findings here support previous results that developmental senescence is not associated with expression of classical inflammatory factors such as *Il1a*, *Il1b* or *Il6*. Altogether, these data now provide a comprehensive and detailed transcriptional profile of the AER and developmental senescence, which we refer to as “DevSen”, and which is provided in Supplementary Table 1.

### Identification of a core, conserved signature across senescent states

A major question relating to developmental senescence is whether there is conservation of a core signature between this and other senescence states. There is overlap of many features and genes, including p21, SA-ß-Gal and some SASP factors. However, whether there is a core molecular signature remains unclear. To address this, we compared the signature of developmental senescence with 18 other senescence datasets (Fig. 3a). We recently performed transcriptome profiling of 18 published in vitro senescence RNA-sequencing datasets, which included many different cell types, with various senescence inducers^21^. This comprised various cell lines (IMR90, MEF, HAEC, HUVEC, Wi38, PMK, Pan02, A549) that had been induced by different triggers such as Irradiation (IR), oncogene-induced senescence (OIS), replicative senescence (RS), and therapy-induced senescence (TIS). We compared the differentially expressed genes (DEGs), either upregulated (Fig. 3b) or downregulated (Extended Data Fig 3a) in the 18 datasets with those from the AER. While there were no genes significantly upregulated in all 19 studies, there were 171 genes upregulated in the AER and in more than half of the 18 datasets, while the most was 4 genes common to 15 out of 18 senescence datasets and the AER. Interestingly, the top 4 common genes (*Arrdc4*, *Cd36*, *Clca2* and *Icam1*) have all previously been linked to senescence (Fig. 3c)^13,22–25^. To look more in more detail, we focused on the top 28 genes, increased in more than 12 datasets and the AER (Fig. 3b, c, Extended Data Fig. 3c). This broader list included additional genes that have been previously linked to senescence, including *ApoE*, *Gpnmb* or *Dpp4*^11,12,26^. In addition, the gene for p21 *(Cdkn1a)* was increased in 14 out of 18 datasets, further supporting the strategy. Altogether, this shows that there is a strong conservation of gene-expression between developmental senescence and other models of cellular senescence.

**Figure 3.**
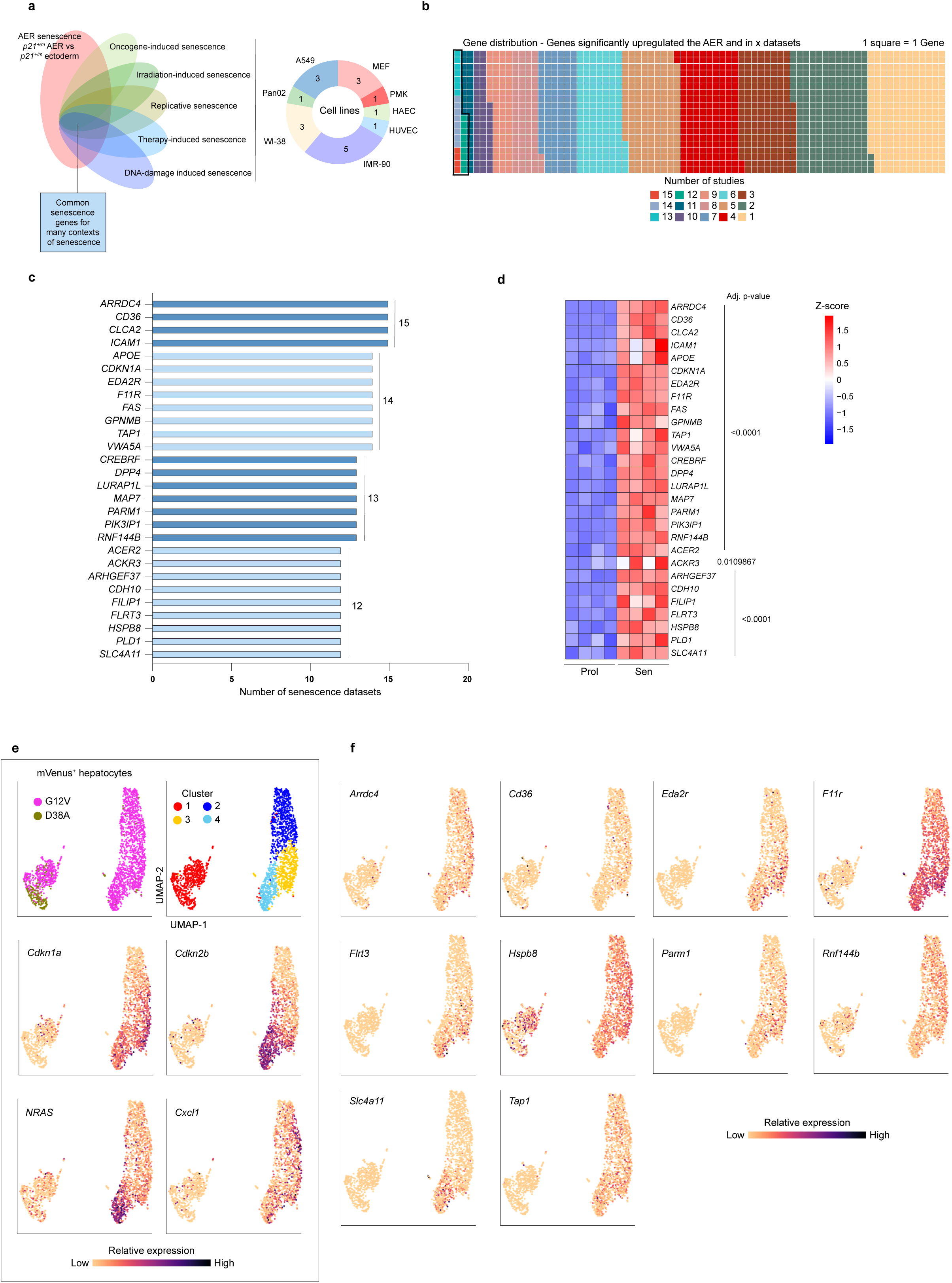
Identification of a conserved set of genes across senescence states. **a**, Left, Venn Diagram showing schematic overlap of the different types of senescence used for the comparative study. Right, pie chart representing the different cell types used in the *in vitro* studies. **b**, Waffle plot showing the distribution of genes significantly up in the AER and in other *in vitro* senescence studies. 1 square = 1 gene, and each color corresponds to the number of studies in which the gene is upregulated. **c**, Top 28 genes obtained by the comparative study and the number of *in vitro* senescence datasets in which they are upregulated. **d**, Heatmap of normalized reads of top 28 genes in proliferating and doxorubicin-induced senescent IMR-90 cells. Row scaling. **e**, UMAP embeddings of single-cell-sequenced hepatocytes colored by experimental condition: cluster, expression of *Nras*, *Cdkn1a*, *Cdkn2a*, *Cxcl1*, and **f**, selected genes of interest.

Our analysis also uncovered genes that were significantly downregulated in developmental senescence and the other models (Extended Data Fig. 3a, b). Interestingly, here we can note the presence of *Psat1*, whose downregulation has been associated with senescence and inhibition of cell proliferation^27^.

Focusing on the commonly up-regulated genes, and to further explore these 28 genes, we next asked if they would be increased in yet another model of senescence that was not included in the meta-study. Here, we induced senescence in IMR90 human fibroblasts using doxorubicin, a well-established model of senescence, and performed bulk RNA-sequencing. Importantly, these cells exhibited classical signs of senescence including enlarged size, SA-ß-Gal positivity and decreased proliferation (Extended Data Fig. 3d, e). From the RNA-sequencing results, we could see increased expression of many senescence-associated genes, including p21, p16, various SASP genes, and decreased expression of Lmnb1, further confirming the senescent status of these cells (Extended Data Fig. 3e). Then, examining the expression of the 28 genes common to developmental senescence and the other models of senescence, we found that they were all significantly increased in doxorubicin induced senescence in IMR90 cells, strongly supporting their association with the senescent state (Fig. 3d).

Given this strong association of these 28 genes with senescence models, we next asked if these genes were associated with another in vivo model of senescence, in this case OIS in liver hepatocytes. A recent study documented senescence patterns in these cells using single-cell RNA-sequencing on oncogenic NRAS^G12V^-expressing hepatocytes^28^. Here, a senescence spectrum was described, identifying 4 clusters of *NRAS*-expressing cells, showing that senescence markers, including both p21 and p15, which are expressed in developmental senescence, were highly expressed in *NRAS*- and SASP-expressing cells (Fig. 3e). We then examined the expression of the 28 genes we identified as being common in developmental senescence and the in vitro senescence datasets. Interestingly, many of these were also increased in the *NRAS*-expressing spectrum, in a similar manner as the senescence markers (Fig. 3e,f, Extended Data Fig. 4). This was particularly evident for F11r, Flrt3, Rnf144b, Eda2r and others, which had highest expression in clusters 3 and 4, overlapping broadly with p21 and p15 expressing cells. To further explore this comparison with in vivo senescence, this same study analyzed the senescence spectrum in an endogenous Kras^G12D^-driven pancreatic tumor model^28^. When we analyzed our 28 genes in this model, again, we could see pronounced shared patterns for p21 and many of the overlapping genes, including Arrdc4, Eda2r, F11R, Flrt3 and others (Extended Data Fig. 5). Together, these results further support the potential involvement of these genes in senescence in vivo.

### Functional screening to identify novel regulators of senescence

Altogether, this strongly supports that developmental senescence and other models of senescence share a significant core gene signature, and hints that there may be additional unknown senescence-regulatory genes or markers common across many states. As a first step to investigate if we could identify novel functional regulators of senescence, we took the approach to knockdown the expression of some of the 28 genes in cells that were induced to undergo senescence, and examine for potential disruption of the program. To achieve this, we treated IMR90 cells with Doxorubicin to induce senescence, and used siRNA to knockdown 10 of the 28 genes, and check for effects on senescence features (Fig. 4a). First, the knockdown efficiency of each gene was examined, which was over 80% decreased in most situations (Extended Data Fig. 6). In each case, cells receiving pools of either a control siRNA or siRNA to one of the genes were imaged for changes in their morphology and stained with phalloidin and DAPI to enable imaging of cell and nuclear sizes (Fig. 4b). In most cases, there was no visible phenotypic differences, either early after knockdown (not shown), or up to 10 days later (Fig. 4b). However, knockdown of ARRDC4 led to increased cell size early after knockdown of the genes, with the appearance resembling earlier entry into senescence. This increase in size was also noticeable at the 10-day stage, with siARRDC4-treated cells much larger and more spread out than siCTRL cells. Most interesting were the cells with knockdown of EDA2R. These cells presented with a completely altered phenotype, looking thin and stretched compared to the senescent controls.

**Figure 4.**
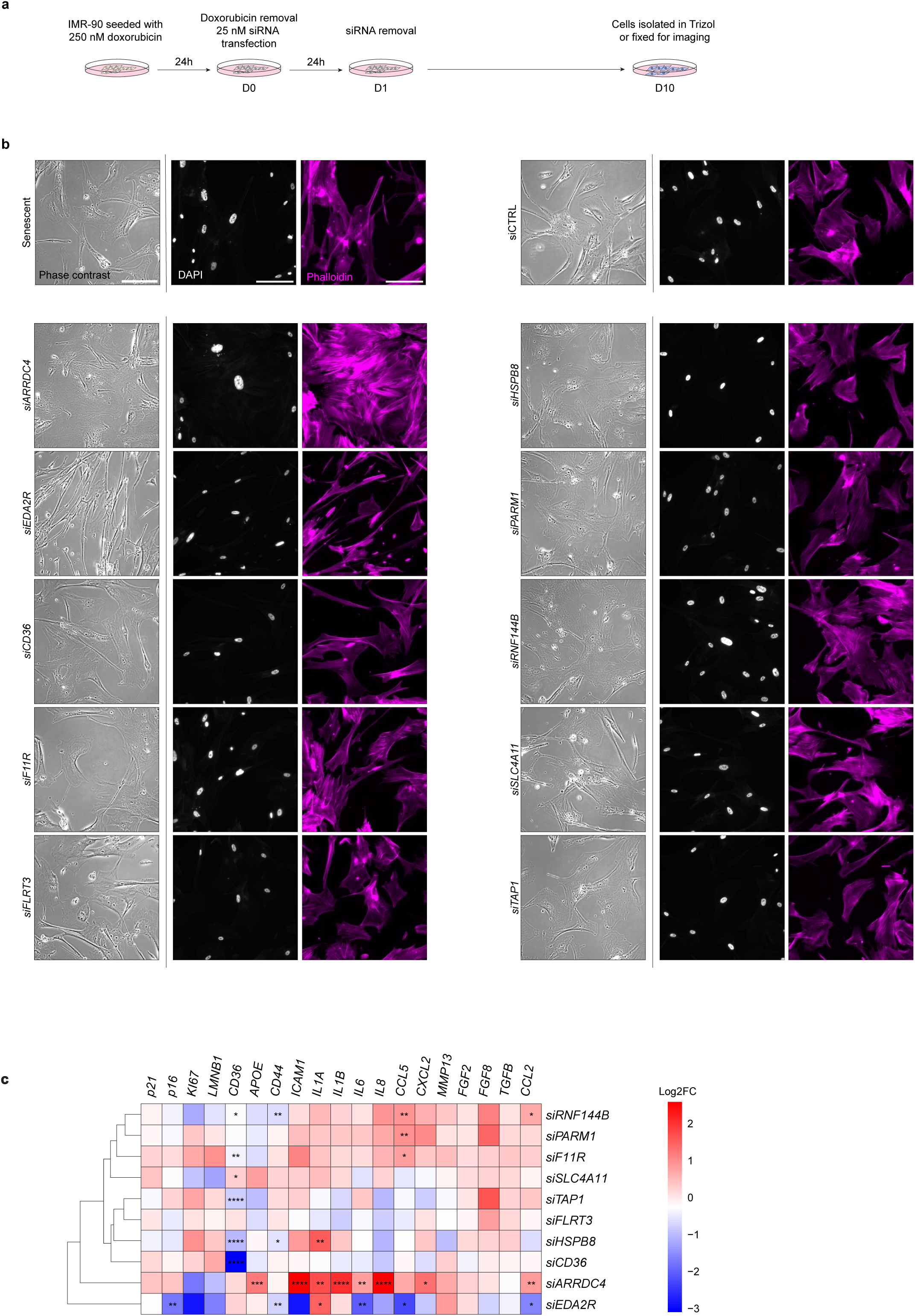
siRNA screening identifies new senescence modifier genes. **a**, Schematic of the siRNA screening experiment protocol to induce senescence and knock-down in IMR-90 cells. **b**, Representative phase contrast, DAPI and Phalloidin pictures of each siRNA, scale bar 200 µm. **c**, Heatmap of log_2_ FC mRNA expression of classical senescent genes measured by RT-qPCR in each siRNA condition, column scaling. Normalized to Rplp0, and to siCTRL of each experiment. n=3, one-way ANOVA test followed by Dunnett’s multiple comparisons, done on each gene and on regular fold change. *: p<0.05, ** : p<0.01, ***: p<0.001, ****: p<0.0001.

To investigate the molecular effect of loss of these genes on senescence, we then performed qPCR to measure the expression of senescence and SASP genes (Fig. 4c). Here the results were more informative and suggested that a number of the genes impacted different aspects of senescence profiles. For example, knockdown of RNF144B and PARM1 genes correlated with increased expression of a number of SASP genes, while knockdown of FLRT3, SLC4A11 and CD36 was associated with decreased expression. Indeed, this latter result is consistent with previous findings positioning CD36 as a SASP activator^25^. Furthermore, knockdown of a number of genes, including TAP1, HSPB8 and F11R resulted in increased expression of KI67, suggesting there may be a functional role in senescence arrest. However, no increased proliferation was noted. Among the most different results, and in agreement with the cellular phenotype, loss of ARRDC4 led to increased expression of p21 and p16, decreased expression of Ki67 and LMNB1, and upregulation of many SASP genes, suggesting that loss of ARRDC4 induces senescence. However, the most striking differences were seen following loss of EDA2R, which resulted in decreased expression of p16, further decreases in KI67 and LMNB1, and a broad downregulation of many SASP genes. In agreement with the striking effect on the senescent cell phenotype, this suggests that EDA2R may be involved in regulating the senescent state.

### Investigating EDA2R as a regulator of senescence features

To examine further a potential role for EDA2R (Ectodysplasin A2 receptor) in senescence, we next investigated its increased expression, this time at multiple stages after senescence induction. Here we could detect increased *EDA2R* as early as 3 days after doxorubicin treatment, similarly to p21 and p16, but even before SASP factors Il6 and Cxcl2 (Fig. 5a). Given the striking phenotype of senescent cells with knockdown of EDA2R 10 days after induction, we next looked at earlier stages after doxorubicin treatment. In agreement with the early induction of EDA2R, the elongate and smaller stretched phenotype became evident as early as 3 days after senescence induction and knockdown (Fig. 5b). It appeared that while the wildtype cells proceeded through the characteristic stages of cell enlargement seen upon senescence induction, cells induced to senesce with EDA2R-knockdown could not proceed through the program, and retained a phenotype of the early stages. Together, these results suggest that increased expression of EDA2R might be an early event in senescence.

**Figure 5.**
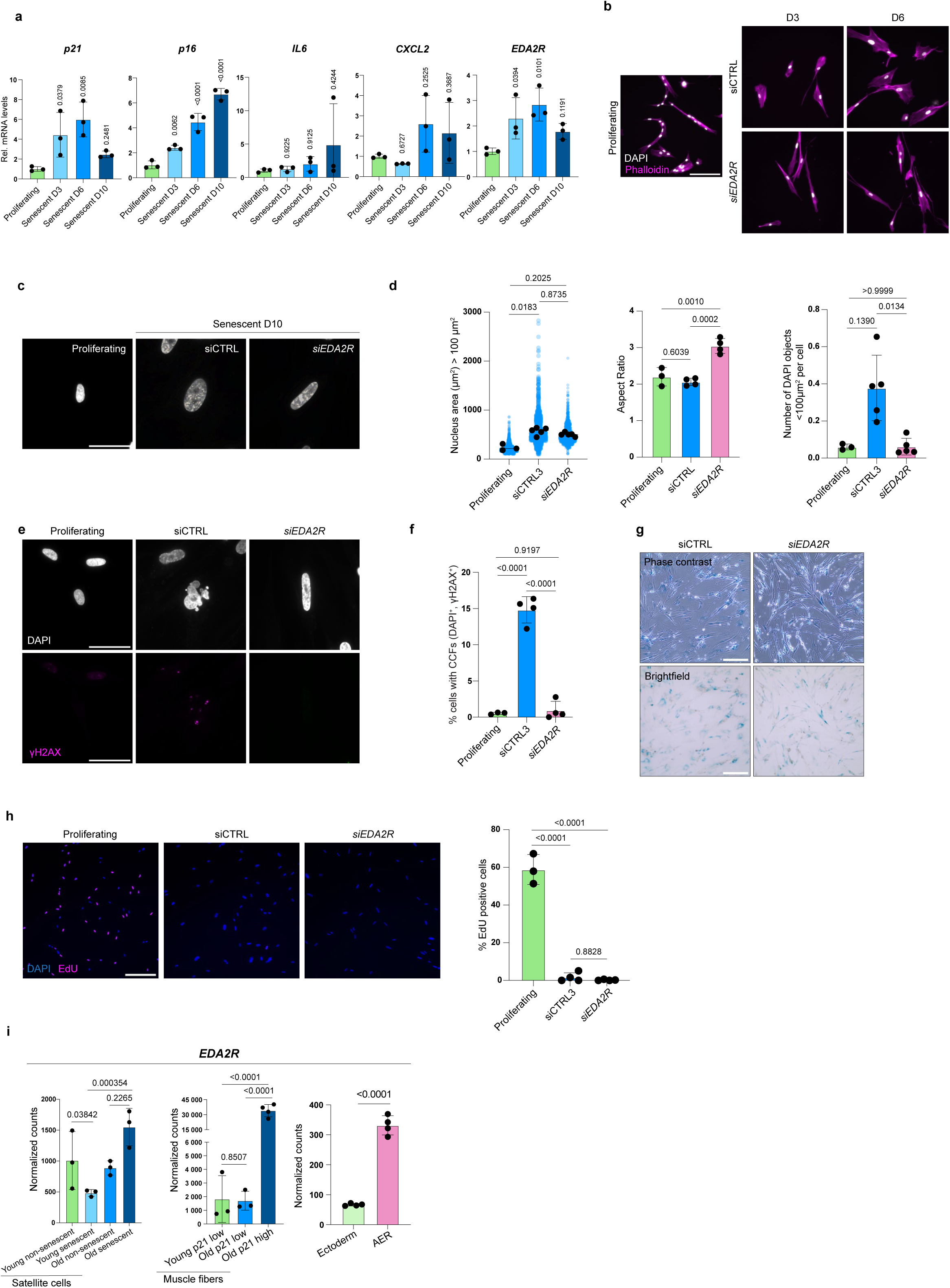
Eda2r regulates many features of the senescent state. **a**, Relative mRNA expression of classical senescence genes and *EDA2R* in proliferating and D3, D6 and D10 doxorubicin-induced senescent IMR-90 cells, n=3. For each gene, one-way ANOVA tests followed by Holm-Šídák’s multiple comparisons tests, compared to average of proliferating samples. **b**, Representative DAPI and Phalloidin pictures of proliferating, Doxorubicin and siCTRL or siEDA2R-treated IMR-90 cells, at D3 and D6 after Doxorubicin removal, scale bar 200 µm. **c**, Representative DAPI pictures of proliferating and senescent IMR-90 cells, treated with siCTRL or siEDA2R. Senescent cells fixed 10 days after doxorubicin removal. Scale bar 50 µm. **d**, Left, Nucleus area of detected DAPI object > 100 µm^2^ in proliferating and D10 senescent treated with siCTRL or siEDA2R in IMR-90 cells. Middle, Aspect ratio (length divided by width) of nuclei of proliferating, siCTRL or siEDA2R-treated D10 senescent IMR-90 cells. At least 60 randomly selected cells were assessed for each replicate. n=3-4, one-way ANOVA followed by Tukey’s multiple comparisons test. Right, Number of detected DAPI objects < 100 µm^2^ per cell in proliferating and D10 senescent IMR-90 cells, treated with siCTRL or siEDA2R. n=3-5, Kruskall-Wallis test followed by Dunn’s multiple comparisons test. **e**, Representative DAPI and γH2AX pictures of proliferating or D10 senescent IMR-90 cells, treated with siCTRL or siEDA2R, scale bar 50 µm. **f**, Percentage of cells with cytoplasmic chromatin fragments (CCFs, DAPI^+^, γH2AX^+^) in proliferating and D10 senescent IMR-90 cells, treated with siCTRL or siEDA2R. At least 60 randomly selected cells were assessed for each replicate. n=4, one-way ANOVA followed by Tukey’s multiple comparisons test. **g**, Representative phase contrast and brightfield pictures of SA-ß-Gal staining of D10 senescent IMR-90 cells treated with siCTRL or siEDA2R, scale bar 300 µm. **h**, Left, Representative DAPI and EdU pictures of proliferating and D10 senescent IMR-90 cells, treated with siCTRL or siEDA2R, scale bar 300 µm. Right, Percentage of EdU positive cells after 24H incorporation, in proliferating and D10 senescent IMR-90 cells, treated with siCTRL or siEDA2R, n=3-4. One-way ANOVA tests followed by Holm-Šídák’s multiple comparisons tests. **i**, Normalized counts of EDA2R in: Left, in young non-senescent, young senescent, old non-senescent and old senescent satellite cells, data extracted from Moiseeva et al., (2023)^13^. Middle, young p21 low, old p21 low and old p21 high muscle fibers, data extracted from Zhang et al., (2022)^31^. Right, the ectoderm and the AER from RNA-seq performed in this study.

We next characterized additional ways in which loss of EDA2R impacted senescence. In addition to the cell shape, the nucleus of cells treated with siEDA2R appeared different to senescent controls (Fig. 5c). As an enlarged nucleus is a hallmark feature of senescent cells, we measured their size and shape. Overall, the nuclei in cells with knockdown of EDA2R were slightly smaller, but were significantly different in their shape, presenting with a more elongate, less round phenotype than siCTRL (Fig. 5d). In this measurement of nuclear area, it also became evident that nuclear objects smaller than 100 µm^2^ present in siCTRL senescent cells were less present in siEDA2R cells. As these were suggestive of cytoplasmic chromatin fragments (CCFs), another feature of senescent nuclei^29^, we assessed their occurrence and found a significant reduction in senescent cells with EDA2R knockdown (Fig. 5e, f). However, when we checked SA-ß-Gal activity, we found no major change in expression levels in senescent cells upon EDA2R knockdown (Fig. 5g). In addition, even though loss of EDA2R significantly impacts many features of senescent cells, these cells remain arrested from proliferation which was confirmed by Edu incorporation analysis and phenotypic observation (Fig. 5h). Altogether, these results uncover a previously undescribed role for EDA2R in mediating key aspects of the senescence program.

Our results uncovered Eda2r as a functional regulator of senescence. Interestingly, increased Eda2r has recently been linked with the aging process^30^, however, no link was made with senescence in this study. Therefore, we analyzed published profiles of senescent cells isolated from muscle, including aged satellite cells^13^ and muscle fibers^31^ (Fig. 5i). Remarkably, in both cases, Eda2r expression was higher in senescent cells, even compared to non-senescent age-matched controls. This was particularly evident in the p21-high subpopulation of muscle fibres, further supporting a link with EDA2R and p21 expression. This is indeed similar to the expression pattern seen in the developing embryonic limb, with high expression of Eda2r in the p21-positive AER compared to the surface ectoderm (Fig. 5i). Altogether, this strongly supports that increased Eda2r is a marker of senescence, both in vitro and in vivo, and at early and late stages of lifespan.

### EDA2R regulates key features of senescence including SASP and protein degradation pathways

Finally, to investigate how EDA2R contributes to the regulation of the senescence program, we performed RNA sequencing on senescent cells with and without EDA2R knockdown (Fig. 6a). Principal component analysis (PCA) revealed marked separation between the two groups, underscoring pronounced transcriptional differences (Fig. 6b). Differential expression analysis identified 4485 significantly upregulated genes and 4325 significantly downregulated genes upon EDA2R loss (Fig. 6c, Extended Data Fig. 7a, Supplementary Table 2). Examination of canonical senescence-associated transcripts reinforced our prior observations (Fig. 4c), showing that knockdown of EDA2R reduced expression of key regulators including *CDKN2A (p16)*, *CDKN2B (p15)*, *CDKN1C (p57)*, *LMNB1*, and *MKI67*, as well as multiple SASP components such as *ICAM1*, *IL1A*, *IL1B*, *CCL2*, and *CXCL8* (Fig 6d).

**Figure 6.**
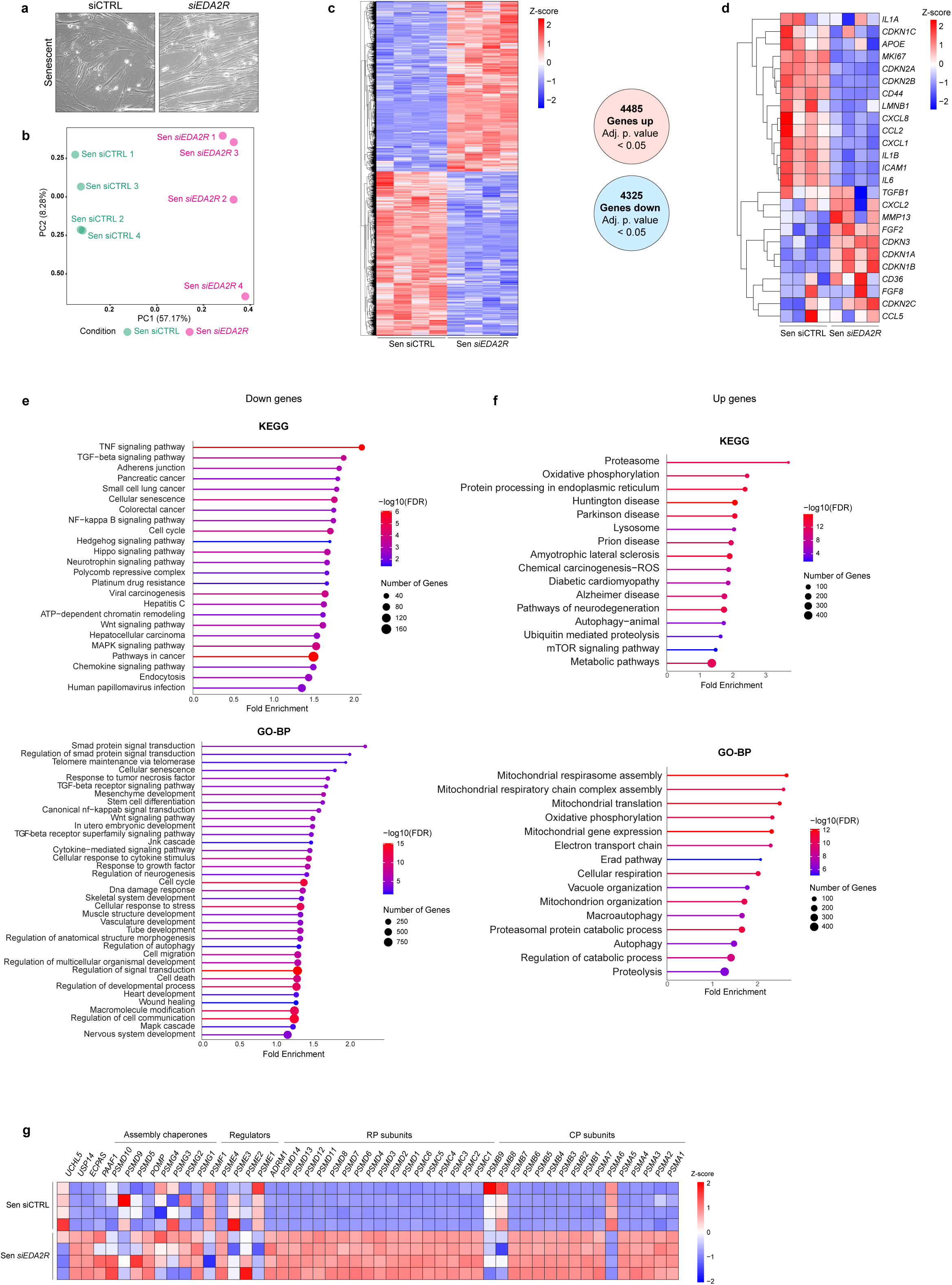
Eda2r regulates secretory and proteostatic features of senescence. **a**, Representative phase contrast pictures of doxorubicin-induced senescent IMR-90 cells treated with siCTRL or siEDA2R for 24h. Cells were collected 6 days after doxorubicin removal for RNA-seq. Scale bar 200 µm. **b**, PCA of senescent siCTRL and senescent siEDA2R samples used for RNA-seq. **c**, Heatmap of normalized reads of differentially expressed genes (DEGs), and number of upregulated and downregulated DEGs, adj. p-value <0.05. Row scaling. **d**, Heatmap of normalized reads of senescent genes in senescent siCTRL and senescent siEDA2R samples. Row scaling. **e**, Selected enriched terms of KEGG and GO-BP analysis for significant downregulated DEGs (Adj. p-value <0.05) between senescent siCTRL and siEDA2R samples. **f**, Selected enriched terms of KEGG and GO-BP analysis for significant upregulated DEGs (Adj. p-value <0.05) between senescent siCTRL and siEDA2R samples. **g**, Heatmap of normalized reads of proteasome genes in senescent siCTRL and senescent siEDA2R samples. Column scaling.

To examine the broader impact of EDA2R depletion, we next conducted Gene Ontology (GO) pathway enrichment analysis. In agreement with the previously known functions of EDA2R, downregulated gene sets included terms linked to *canonical NF-κB signal transduction* and *Jnk cascade*. Importantly, these are both key senescence pathways, regulating SASP activation and cytoplasmic chromatin fragment (CCF) formation respectively, further supporting a role for EDA2R in sustaining these senescence-associated features (Fig. 6e, Extended Data Fig. 7b, d, e). In addition, other senescence-associated downregulated terms included *cellular senescence*, *cytokine-mediated signaling pathway* and *DNA damage response.* Furthermore, genes associated with developmental signaling pathways, such as TGF-beta, Hedgehog, Hippo, Wnt and MAPK signaling were also diminished, suggesting that EDA2R may govern a broader developmental secretory repertoire in senescent cells beyond classical SASP factors (Supplementary Table 3).

Conversely, analysis of the upregulated transcriptome uncovered unexpected results (Fig. 6f, Extended Data Fig. 7c). Interestingly, KEGG analysis identified gene sets linked to signatures of multiple neurodegenerative disorders, including *Prion*, *Huntington*, *Parkinson’s*, and *Alzheimer’s disease,* in addition to A*myotrophic Lateral Sclerosis*. Most prominent among these signatures however was *proteasome*, in addition to other protein degradation pathways *lysosome* and *autophagy*. As impaired protein degradation is a key feature of senescence and neurodegenerative diseases^32,33^, this suggests that loss of EDA2R restores some of these functions. Indeed, closer inspection of the proteasomal pathway revealed a significant induction of expression of proteasome subunit genes, whose decline is a key feature of senescence (Fig. 6g).

Taken together, these data suggest that elevated EDA2R expression in senescent cells drives the activation of SASP and CCF-associated signaling as well as additional secretory pathways, while concomitantly repressing proteasomal function and protein degradation. This dual role positions EDA2R as a central regulator of both the pro-inflammatory and degradative arms of the senescence program.

## DISCUSSION

Cellular senescence is implicated in tumor suppression, aging, and various diseases, but also plays beneficial roles in embryonic development, wound repair, and tissue regeneration^2–4,18^. These opposing functions have led to the hypothesis that senescence is a fundamentally beneficial process that becomes dysregulated in pathological and aged contexts. This interpretation therefore implies the existence of a shared molecular program common to senescence across diverse states. However, direct comparisons between senescent cells in beneficial versus detrimental contexts have been limited, leaving it unclear whether such a conserved core signature exists.

We have previously shown that the apical ectodermal ridge (AER), a key developmental signaling center, displays hallmarks of senescence, including p21 expression, SA-β-Gal activity, reduced proliferation, and a pronounced secretory phenotype^15^. Building on this, we aimed to transcriptionally profile this population of developmentally senescent cells in greater detail. Our rationale was that while senescence markers like p21 and SA-β-Gal are transiently expressed during embryonic senescence, they are persistently activated in aging and pathological contexts^7,34^, suggesting that other genes may follow similar temporal dynamics.

To enable specific isolation of the senescent AER, we generated a new reporter mouse model that labels p21-expressing cells. This model was rigorously validated both *in vitro* and *in vivo*, confirming that while the targeting disrupts the endogenous *Cdkn1a* locus, heterozygous mice are phenotypically comparable to wild-type. This reporter line now provides a versatile tool for studying p21 and senescence across developmental and adult contexts, enabling precise cell identification, isolation, tracing, and targeted ablation. Transcriptomic profiling confirmed enrichment of established AER markers, including *Fgf8*, *Fgf4*, and *Cd44*, validating the specificity of the isolated population. While we previously profiled the AER using microarray-based approaches, the new model allowed isolation of a purer population of senescent cells, enabling deeper transcriptomic analysis via RNA sequencing. The resulting gene signature, which we refer to as “DevSen,” now serves as a valuable resource for investigating the molecular landscape of developmental senescence and for comparing this to senescence in other biological contexts.

A persistent challenge in the senescence field is the lack of robust markers to reliably identify senescent cells across contexts. With this in mind, we compared the “DevSen” profile of the senescent AER with a range of senescence models, aiming to identify genes consistently upregulated across diverse contexts. By focusing on genes that were elevated both in developmental senescence and in other stress-induced models, we identified a shared set of transcripts that define a conserved senescence signature. Notably, this analysis recovered many genes previously implicated in senescence, either as markers or functional mediators. Indeed, the most consistently upregulated were *Arrdc4*, *Cd36*, *Clca2*, and *Icam1*, all of which have prior associations with senescence^22–25^. Additional genes such as *Dpp4*, *Gpnmb*, and *Apoe*, alongside *Cdkn1a* (p21)^7,11,12,26^, further validated the approach and underscored the biological relevance of the conserved set. Several of these proteins, including Dpp4, have already been proposed as surface markers of senescence, and our data provide independent support for their utility. Importantly, this conserved signature was not restricted to in vitro models, with many of these genes also elevated in oncogene-induced senescence (OIS) in hepatocytes and pancreatic cells *in vivo*, reinforcing their broader applicability. These findings validate our comparative strategy and demonstrate the existence of a core gene-set shared by senescent cells across developmental, physiological, and pathological settings.

To identify potential new mediators of senescence, we performed a focused functional screen targeting 10 of the top 28 genes from our conserved signature. Using gene-knockdown approaches, we assessed effects on both senescence-associated gene expression and cellular phenotype. This screen revealed several candidates that may contribute to discrete aspects of the senescence program, though their roles may be context-dependent and will require further investigation. For example, knockdown of *Tap1*, *Parm1*, *Hspb8*, and *F11r* led to modest increases in Ki67 expression, suggesting partial release from cell cycle arrest. In the case of *Flrt3*, a gene known to be essential for AER cell survival^35^, knockdown in senescent cells did not induce notable cell death, but was associated with a trend towards reduced p16 and SASP expression, alongside elevated Ki67. These observations raise the possibility that some of these genes may contribute to senescence maintenance or survival, though more stable knockdown models, and *in vivo* studies, may be needed to clarify their roles. Among all candidates, knockdown of *Arrdc4* produced one of the most pronounced phenotypes. Cells became notably larger compared to senescing controls, which was accompanied by increased expression of p16 and SASP genes, and decreased Ki67 and Lmnb1, hallmarks of a reinforced or accelerated senescence. Interestingly, many of the top-ranked genes in our signature are known NF-κB targets, including *Tap1*, *F11r*, *Apoe*, *Cdkn1a*, *Fas*, and *Tnfsf15*. Others, such as *Eda2r* and *Cd36*, function upstream of NF-κB signaling. Given NF-κB’s central role in regulating the SASP, these findings reinforce that NF-κB activity may represent a conserved regulatory axis underpinning senescence across diverse biological contexts, and implicate additional targets in this role.

The selective elimination or modulation of senescent cells in aged or damaged tissues has been shown to confer therapeutic benefits in multiple preclinical models, driving interest in senescence-targeting strategies^9,10^. A central concern, however, is that interventions should spare beneficial, transient senescence, while selectively targeting detrimental or chronic senescent states. Our findings complicate this distinction, as several genes already used as targets in senolytic or senostatic strategies, including Cd36, Dpp4, and Gpnmb^11–13^, are part of the conserved core signature shared with developmental senescence. Reinforcing this point, we have previously shown that p21 disruption by senotherapeutics can promote liver regeneration^36^, while other studies have reported that clearance of p21-expressing cells in aged tissues improves tissue function^7,37^. These observations suggest that even core components of a beneficial senescence program, now potentially including Eda2r, may become aberrantly or persistently activated in pathological settings, and that their transient inhibition could be therapeutically advantageous. Ultimately, our work also opens the possibility of identifying genes that are unique to damage- or stress-induced senescence, which could serve as more selective targets to refine therapeutic approaches and reduce off-target effects.

Finally, we focused our analysis on Eda2r. Among the candidates tested, Eda2r emerged as a particularly compelling regulator, with its loss resulting in reduced expression of multiple core senescence and SASP features. Notably, Eda2r was recently implicated in cancer cachexia^38^ and was identified as a biomarker of aging^30^. However, these studies did not directly explore a functional link between Eda2r and cellular senescence. To investigate this connection, we analyzed transcriptomic data from senescent cells isolated from aged tissues and found that Eda2r expression was significantly elevated compared to non-senescent, age-matched controls. This suggests that increased Eda2r levels in aging are likely driven by its upregulation specifically in senescent cells.

The effect of Eda2r knockdown on the senescence phenotype was notably pronounced. Interestingly, its loss did not alter p21 expression or trigger senescence bypass through increased proliferation. Instead, Eda2r knockdown selectively disrupted multiple hallmark features of the senescent state, including elevated p16 expression, increased nuclear size, cytoplasmic chromatin fragment (CCF) formation, and the SASP. These results suggest that Eda2r is not essential for initiating cell cycle arrest, but is required for the full establishment or maintenance of the senescence program. This is consistent with previous reports implicating EDA2R in regulating NF-κB and JNK signaling, pathways central to SASP production and CCF formation, respectively^39^. Interestingly also was the finding that senescent cells without Eda2r had significantly reduced expression of many developmental and senescence-associated signaling pathways, including Tgf, Tnf, Smad, Mapk and others. This also fits with our recent study^21^ suggesting senescent cells have a core developmental signaling component. Finally, and somewhat unexpectedly, Eda2r knockdown in senescent cells led to a significant increase in expression of genes associated with protein degradation, including the proteasome and autophagy, in addition to signatures associated with mitochondrial function. These results raise the possibility that Eda2r contributes to senescence maintenance by reinforcing proteostatic and mitochondrial stress, and uncover Eda2r as a new and potentially actionable regulator of senescence.

Overall, this study reveals the existence of a conserved core gene signature shared across diverse senescent contexts, including developmental senescence, oncogene-induced senescence (OIS), replicative senescence (RS), and therapy-induced senescence (TIS). Much like p21, these genes are expressed in specific developmental contexts but become aberrantly or persistently upregulated in pathological senescence. While the precise roles of individual genes within this conserved program remain to be fully elucidated, our functional and mechanistic analyses highlight Eda2r as a novel regulator of senescence maintenance. These findings not only provide a valuable framework for dissecting the molecular architecture of senescence but also point toward new avenues for potential therapeutic targeting in aging, cancer, and regenerative contexts.

## METHODS

### Animals models and genotyping

C57Bl/6J (WT), p21-KO and p21-mCherry-CreERT2 mice were bred and maintained in temperature and humidity-controlled environment at the animal facility Institut Clinique de la Souris (ICS) of the IGBMC with a 12-hour light/dark cycle, in compliance with French and European Union regulations on the use of laboratory animals for research. The p21-KO mice were on a C57Bl/6J genetic background, while the p21-mCherry-CreERT2 were 50:50 C57Bl/6J:C57Bl/6N. The genotyping was performed on a tip of the tail using Mouse Direct PCR Kit (Bimake B40013) according to the manufacturers protocol. For PCR genotyping of p21-mCherryCreERT2 mice, the following primers were used: p21CherryCreERT2_fw: 5’-CCCTTAGACTCTGGGGAATGATGTCC -3’; p21CherryCreERT2_rv: 5’-CCCTCTTCATCCGGTAAAGCAACTCG -3’ and p21CherryCreERT2_wt: 5’-GGGCCCTACCGTCCTACTAA -3’. Combination p21CherryCreERT2_fw/p21CherryCreERT2_rv highlights presence of transgene allele at 152 bp and combination p21CherryCreERT2_fw/ p21CherryCreERT2_wt highlights presence of WT allele at 357 bp.

### Generation of p21-mCherry-CreERT2 model

To generate the p21-mCherry-CreERT2 knock-in mice, genomic DNA encompassing the p21 locus was amplified from BAC RP23-73N16 using high-fidelity PCR. The resulting DNA fragments were assembled with a LF2A-mCherry-T2A-CreERT2 cassette and were inserted in-frame immediately upstream of the endogenous stop codon into the targeting vector. This targeting vector was co-electroporated as a circular plasmid along with a CRISPR/Cas9 expression plasmid (pX330) into C57BL/6NCrl (S3) embryonic stem (ES) cells derived in-house. The CRISPR plasmid encoded wild-type *SpCas9* and a single guide RNA (gR88; sequence: gggctcccgtgggcacttca) designed to induce a double-strand break at the target site, thereby enhancing homologous recombination efficiency. G418-resistant ES cell colonies were screened for correct targeting by PCR and confirmed by Southern blot hybridization, protocols^40^. Two correctly targeted clones with normal karyotypes were microinjected into BALB/cN blastocysts, which were subsequently transferred into pseudopregnant recipient females. Chimeric offspring were bred to C57BL/6N Flp deleter mice^41^ to excise the floxed selection cassette. Germline-transmitting progeny carrying the desired allele—free of both the selection marker and the *Flp* transgene—were identified by genotyping.

### Isolation and culture of Mouse Dermal Fibroblasts

Primary mouse dermal fibroblasts were isolated from the skin of C57Bl/6J (WT), p21-KO and p21-mCherry-CreERT2 pups, 0–1-day old. Newborn pups were euthanized by anesthetic injections of firstly xylazine (Rompun®, 1/10 in NaCl) and secondly pentobarbital sodium and phenytoin sodium (Euthasol®, 1/20 in NaCl). Skins were dissected and washed with PBS 1X before being incubated overnight in 2.5 mg/mL dispase II (Sigma 4942078001) epidermis up at 4°C. The next morning, the dermis was separated from the epidermis, minced, and incubated in filtered 0.25% collagenase A (Sigma-Aldrich 10103586001) in PBS 1X at 37°C for 30 minutes. Then DNase I (Roche 11284932001) at 10 mg/mL was added, and skins were incubated at 37°C for 5 min. Culture media (DMEM 4.5g/L Gibco 41966 glucose with glutamax-I, 10% FCS (Fetal Calf Serum), NEAA (Non-Essential Amino Acid), gentamycine 40µg/mL and 100µm ß-mercaptoethanol) was added and dissociated cells were filtered through 70 µm cell strainer (Corning 352350), collected by centrifugation and seeded in 15 cm petri dishes. MDFs obtained were then cultured at 37°C in 5% CO2 in DMEM (Gibco 41966) 4.5g/L glucose supplemented with glutamax-I, 10% FCS, NEAA, gentamycine 40µg/mL and 100µm ß-mercaptoethanol. The number of replicates indicated in the figure legends corresponds to the number of batches of isolated MDFs, each from different embryos. For each experiment, the isolated MDFs come from at least three different litters.

### Culture of IMR-90

IMR-90 were cultured in DMEM (Gibco 41966) supplemented with 10% FCS and 1% penicillin-streptomycin at 37°C in 5% CO2.

### Senescence induction

Radiation-induced senescence in MDFs was induced by exposing the cells to a dose of 10Gy of X-rays using a CellRad Precision X-ray irradiator. Settings used were 130 kV and 5 mA at tray-position one, for a dose rate of 0.92 Gy/min. For therapy-induced senescence in IMR-90, cells were treated with 250 nM doxorubicin for 24H^42^. Cells were then washed and cultured in normal culture medium described above.

### SA-ß-Galactosidase staining

IMR-90 stained for SA-ß-Galactosidase activity were washed and fixed in 0.5% glutaraldehyde (Sigma G7651) for 15 minutes at RT. Fixed cells were washed twice with PBS 1X containing 1 mM MgCl2 at pH6 and kept at 4°C until SA-ß-Gal experiment (no more than a week). Cells were stained in X-Gal solution consisting of 1 mg/mL X-Gal (Biosynth AG), 5 mM K3Fe(CN)6 and 5 mM K4Fe(CN)6 3H2O in PBS1X 1 mM MgCl2 at PH=6 for 8-12 hours. Cells were washed with PBS 1X and imaged with brightfield microscope (EVOS XL Core). Quantification of SA-ß-Gal positive cells was done manually with Fiji software by counting cells in 3-4 pictures (field of view 1000 µm) by condition. Embryos stained for SA-ß-Galactosidase activity were dissected into PBS 1X and fixed in 0.5% glutaraldehyde (Sigma G7651) overnight at 4°C with gentle agitation. Fixed embryos were then washed 3 times for 10 minutes with PBS 1X mM MgCl2 at PH=5.5 and kept at 4°C until SA-ß-Gal experiment (no more than a week). Embryos were stained in X-Gal solution consisting of 1 mg/mL X-Gal (Biosynth AG), 5 mM K3Fe(CN)6 and 5 mM K4Fe(CN)6 3H2O in PBS 1X mM MgCl2 at PH5.5 for 3-8 hours until AER or otic vesicles were positive. Embryos were washed with PBS 1X and imaged with macroscope Leica MacroFluo Z16 APO.

### Focus Formation Assay

Cells were seeded at 2,500, 5,000 and 10,000 cells per wells in 6-wells plates and were grown in culture for 10 days, with frequent medium change. After 10 days, cells were washed with PBS 1X, then fixed in 4% PFA for 15 min at RT. After washing with PBS 1X, staining was conducted by incubating the cells in solution of 0.5% crystal violet (Sigma-Aldrich V5265) and 20% methanol for 20 min at RT by rocking gently. Stained plates were then washed gently with water 4 times, and air-dried. The photos of the plates were taken using a conventional scanner.

### EdU incorporation

EdU incorporation was done on MDFs using Click-iT® EdU Alexa Fluor® 488 Imaging Kit (ThermoFisher C10337) according to manufacturer’s protocol. Briefly, cells were incubated in medium containing final concentration of 10 µM EdU for 2H15 for MDFs and 24H for IMR-90 at 37°C. Cells were then fixed with 4% paraformaldehyde (PFA; Electron Microscopy Sciences, 15710, not included in EdU kit) in PBS 1X, washed with 3% BSA (Sigma A8806, not included in EdU kit), and permeabilized with 0.5% Triton® X-100 (Sigma, T8787, not included in EdU kit). Click-iT® reaction was then performed according to kit’s protocol for 30 min at RT. Nuclei were stained with DAPI at 0.1 µg/mL (Invitrogen D1306) for 2 minutes. Cells were then imaged on Zeiss Axio Observer Z1 and EdU-positive cells were quantified using Cell Profiler, using 15 pictures per conditions and per experiment, field of view 1000 µm. The percentage of EdU-positive cells was obtained by dividing the number of EdU-positive cells by the number of DAPI-positive cells.

### RNA isolation for RT-qPCR

RNA from proliferating and irradiated MDFs was extracted by using NucleoSpin® RNA Plus kit (Macherey-Nagel 740984.250) according to the RNA extraction protocol. RNA from IMR-90 was isolated by Trizol (Thermofisher scientific 15596026) – chloroform protocol by following manufacturer’s protocol for cells in culture. RNA from ectodermal jackets was isolated by Trizol-chloroform protocol following the same steps described above used for samples sent for RNA-sequencing. RT-qPCR on MDFs and IMR-90 was done by using qScript cDNA Super Mix (VWR INTERNATIONAL SAS 733-1177) for the retro transcription and SYBR Green I Master (Roche 04887352001) for qPCR reaction, by following manufacturer’s instructions. As EJs samples provided very little RNA, RT-qPCR was performed in One-Step by using Luna Universal One-Step RT-qPCR Kit (NEB E3005L) by following the kit’s protocol. Reactions for both RT-qPCR and One-Step-qPCR were done in thermocycler LightCycler 480 (Roche). Relative levels of expression were normalized to Rplp0 or ß-actin (indicated in the legends of figures), and to the mean CT values of control samples. Gene-specific primer sequences indicated in Supplementary Table 4.

### Immunofluorescence staining

In the various conditions, cells were fixed in 4% PFA for 15 min at RT and then washed 3 times with PBS 1X. Cells were then permeabilized for 10 min in 0.1% Triton-X in PBS 1X, washed in PBS 1X thrice and then blocked with 10% goat serum, 0.1% Tween in PBS 1X for at least one hour at RT. Cells were then incubated with primary antibody (in 1% goat serum, 0.1% tween in PBS 1X) for at least 2 hours at RT, followed by 3 washing with PBS 1X. Secondary antibody (in 1% goat serum, 0.1% tween in PBS 1X) solution was then added for 1 hour at RT, before being washed 3 times with PBS 1X. If necessary, phalloidin was added for 30 min according to manufacturer’s instruction (ThermoFisher, A12380). Nuclei were stained with DAPI 1 µg/mL for 2-3 min at RT. Cells were imaged with Zeiss Axio Observer Z1 or upright widefield microscope Leica DM6B-Z. Primary antibodies were used at the following concentrations: anti-p21 (Hugo291, undiluted supernatant provided by CNIO), anti-RFP (1/500, Rockland, 600-401-379), anti-γH2AX (1/1000, Merck – 05-636).

### Whole-mount in situ hybridization

RNA probes were prepared by in vitro transcription using the Digoxigenin-RNA labeling mix (Roche). Template plasmid for p21 was kindly provided by Dr. P. Dollé, IGBMC. Whole-mount in situ hybridization were performed as previously published^43^.

### Protein isolation

Cells were washed with PBS 1X, dissociated with Trypsin 0.05% and centrifuged to obtain a cell pellet. After the supernatant was removed, the pellet was resuspended in lysis buffer (RIPA buffer-Bio Basic RB4476, PMSF-Sigma, DTT -Euromedex and PIC-Roche 04693132001) and incubated in this buffer for 20 min at 4°C with shaking. Samples were then sonicated (Bioruptor®Twin, Diagenode) and centrifuged for 10 min at 4°C. Supernatants were transferred to new tubes and quantified using Bradford reagent (BioRad 5000112) according to manufacturer’s instructions, and using a standard range prepared with BSA. Protein concentration was determined by measuring absorbance at 750 nm. Embryos dissected for Western Blot were snap frozen in liquid nitrogen and kept at -80°C until the day of experiment. The day of extraction, embryos were incubated with lysis buffer described earlier and were dissociated with beads in a shaker. Embryos were then vortexed and centrifuged at 11,000g for 5 min at 4°C. Supernatants were transferred to new tubes and quantification was conducted as described for cells.

### SDS Page and Western Blotting

Protein samples were prepared for SDS-Page by adding Laemmli buffer (BioRad 1610747) and 1/10 ß-mercaptoethanol, and were incubated 10 min at 70°C. Protein content was then separated on 15% pre-cast polyacrylamide (Biorad 1610140) resolving gel by running the gel 1-2H at 20-30 mA before being transferred to a 0.45 µm nitrocellulose membrane (GE Healthcare 10600002) by running the blot at 100V for 2H at 4°C. Membranes were stained with Ponceau 0.1% in 5% acetic acid to check the transfer. Membranes were then blocked with TBS-tween 0.05% (25 nM Tris-HCl, PH=7.5, 150 nM NaCl, 0.05% tween) and 5% milk for 1 hour at RT with shaking. Blocked membranes were incubated with primary antibody overnight with shaking at 4°C in TBS-Tween and 2.5% milk. The next day, after 3 washes in TBS-tween, the membranes were incubated with secondary antibodies coupled to HRP (diluted in TBS-tween + 2.5% milk) for 1 hour at RT. After washing in TBS-tween, membranes were developed using regular enhanced chemiluminescence (ECL), Pico or Femto chemiluminescent substrate (respectively ThermoFisher 32209, ThermoFisher 34577 and ThermoFisher 34095). Primary antibodies were used at the following concentrations: anti-p21 (1/500, BD Pharmingen 556431), anti-RFP (1/5000, ROCKLAND, 600-401-379) and anti-α-Tubulin (1/5000, Sigma-Aldrich T6199).

### mCherry intensity measurement

Living proliferating or irradiated MDFs from *p21^+/m^*mice were photographed using the Zeiss Axio Observer Z1 every day for 4 days after irradiation. Quantification was performed on 15-20 pictures (of 211.30 µm field of view) per condition and experiment, and mCherry signal was measured in Fiji using StarDist (Schmidt et al., 2018). In total, between 20 to 298 cells were analyzed for their mCherry intensity in each condition and experiment. Living *p21^+/m^* or *p21^m/m^* embryos were photographed with Leica MacroFluo Z16 APO.

### Isolation of ectodermal jackets for One-Step qPCR

The dissociation protocol was based on Reinhardt et al.’s method to obtain mesenchymal progenitors and was adapted and optimized for our conditions^44^. Forelimb buds were dissected from WT C57Bl/6J, heterozygous or homozygous p21-mCherry-CreERT2 E11.5 embryos in cold PBS 1X. Dissected limbs were pooled (around 12-18 forelimbs coming from one litter) and digested at 4°C in 2% trypsin (Gibco 15090046) for 5 min with gentle shaking. Digestion was stopped in 10% FCS in PBS 1X, and digested limbs were strongly vortexed until ectoderm (ectodermal jackets, EJ) and mesenchyme separated. Ectodermal jackets were retrieved in another clean petri dish and mesenchyme samples were trashed. EJs were then collected in Trizol for RNA extraction.

### Fluorescence-Activated Cell Sorting (FACS) isolation of AER cells

After isolation, EJs were then digested in 0.05% trypsin at RT with agitation, and physically dissociated every 5-10 minutes with p200 until total tissue dissociation. Reaction was stopped by adding 10% FCS PBS 1X and centrifuged for 5 minutes. Pellet was rinsed with PBS 1X and centrifuged again before being resuspend in Pre-Sort Buffer (BD Biosciences-563503). Single-cell suspension was passed through a 50 µm filter and kept on ice before going through sorter. Sorting was done with the help of the Flow Cytometry platform of the IGBMC. Gating was done according to mCherry fluorescence with the help of negative controls (obtained from C57Bl/6J dissected limb). Example of gating in Extended Data Fig. 2g. Sorted cells were retrieved in Pre-Sort Buffer, centrifuged at 4°C for 5 min, rinsed and centrifuged again, before adding 800 µL Trizol (Thermofisher scientific 15596026). Cells in Trizol were then snap frozen in liquid nitrogen and kept at -80°C before proceeding with RNA extraction.

### RNA isolation for AER RNA-sequencing

Cells obtained after sorting in Trizol were thawed on ice and homogenized by vortexing before being incubating 6 min at RT. 160µL of chloroform was added and vortexed for 15 seconds, before incubation 3 min at RT. Samples were centrifuged at 12,000g for 15 min at 4°C. Aqueous phase was transferred to a new tube containing 160µL chloroform. Vortexing and centrifugation was performed again, and aqueous phase was transferred to a new empty tube. 1µL of glycogen 20 µg/µL (ThermoFisher, R0551) was added and samples were homogenized before adding 400µL isopropanol. Homogenized samples were incubated 10 min at RT before being centrifuged at 12,000g for 10 min at 4°C. Supernatants were removed and pellets were washed once with 100% EtOH and twice with 75% EtOH. Pellets were dried at RT and resuspended in RNase-free water. RNA concentration was assessed using Nanodrop 2000c (Thermofisher), and samples were kept at -80°C until the day the samples were donated to the GenomEast platform.

### RNA-sequencing and analysis

Library preparation was performed at the GenomEast platform at the Institute of Genetics and Molecular and Cellular Biology using Illumina Stranded mRNA Prep Ligation - Reference Guide - PN 1000000124518. RNA-Seq libraries were generated according to manufacturer’s instructions from 40 ng of total RNA using the Illumina Stranded mRNA Prep, Ligation kit and IDT for Illumina RNA UD Indexes Ligation (Illumina, San Diego, USA). Libraries were sequenced on an Illumina NextSeq 2000 sequencer as single read 50 base reads. Reads were preprocessed to remove adapter, polyA and low-quality sequences. After this preprocessing, reads shorter than 40 bases were discarded for further analysis. These preprocessing steps were performed using cutadapt version 4.2^45^. Reads were mapped to rRNA sequences of RefSeq and GenBank databases using bowtie version 2.2.8^46^. Reads mapping to rRNA sequences were removed for further analysis. Remaining reads were mapped onto the GRCm39 assembly2 of Mus musculus genome using STAR version 2.7.10b^47^. Gene expression quantification was performed from uniquely aligned reads using htseq-count version 0.13.5, with annotations from Ensembl version 108 and “union” mode3^48^. Only non-ambiguously assigned reads to a gene have been retained for further analyses. Read counts have been normalized across samples with the median-of-ratios method proposed by Anders et al, to make these counts comparable between samples^48^. Hierarchical clustering for SERE coefficient heatmap was performed using the Unweighted Pair Group Method with Arithmetic mean (UPGMA) algorithm. Comparisons of interest were performed by the GenomEast platform using the test for differential expression proposed by Love et al. (2014)^49^ implemented in the Bioconductor package DESeq2 version 1.34.0 and p-value adjustment method of Benjamini and Hochberg (1995)^50^. These genes with an adjusted p-value lower than 0.05 and absolute value of log2 Fold-Change greater than 1 were considered significantly differentially expressed. Heatmaps of normalized reads were created using Pheatmap (1.0.12) package in Rstudio.

### In vivo single-cell analysis

Data generation is as described in Chan et al^28^. For analysis here, resulting reads were aligned using the CellRanger pipeline to the mm10 genome assembly. Demultiplexing based on expression of hashtag oligos was performed using the CITE-seq-Count command, with no mismatches allowed. As all conditions to be compared were pooled into the same experimental run, direct analysis could be performed without the need for integration or batch correction. After quality-control filtering to remove low-quality sequenced cells, all downstream analysis was performed using the Seurat63,64 implementation in R.

### Pathway analysis

Over Representation Analysis (ORA) was performed with clusterProfiler^51^, using the Kyoto Encyclopedia of Genes and Genomes (KEGG) database or with gprofiler2 (version 0.2.3, gprofiler version e113 eg59 p19 f6a03c19)^52^, using Gene Ontology (GO). For each dataset, genes displaying a base mean (average counts across all samples and replicates) of less than 25 bases were excluded prior to analysis. The remaining significant genes with an adjusted p-value below 0.05 and when expressed more or less than +/- 1.5-fold in the AER, were used for the ORA.

Regarding ORA use for the siEDA2R RNA-seq, it was performed using ShinyGO63 (Ge SX, Jung D & Yao R, Bioinformatics 36:2628–2629, 2020) Significantly differentially expressed genes were determined by an adjusted p-value <0.05. Fold enrichment corresponds to the ratio of the percentage of differentially expressed genes present in a pathway term in the complete list of differentially expressed genes, divided by the percentage of genes for a pathway term in all genes detected by RNA-seq.

### Eighteen Senescence dataset analysis

The transcriptome analysis of the 18 senescence datasets presented in this paper was conducted and the methods described in the recent paper^21^.

### Knock-down using siRNA on Doxorubicin-treated IMR-90

siRNA non-targeting control (D-001810-03-05) and against human ARRDC4 (L-019366-02-0005), CD36 (L-010206-00-0005), EDA2R (L-008044-00-0005), F11R (L-005053-00-0005), FLRT3 (L-010090-01-0005), HSPB8 (L-005006-00-0005), PARM1 (L-020227-01-0005), RNF144B (L-025119-00-0005), SLC4A11 (L-007581-02-0005) and TAP1 (L-007634-00-0005) were ordered at Dharmacon and were resuspended according to manufacturer’s instructions. Cells were seeded at 100,000 cells/well in 6-wells plate with 250 nM doxorubicin (Sigma, 44583-1MG) and incubated at 37°C and 5% CO2 for 24 hours. After doxorubicin removal, cells were washed with antibiotic-free medium containing serum. Transfection solution of siRNA and Lipofectamin™ RNAiMAX (ThermoFisher 13778075) was prepared in serum-free and antibiotic-free Opti-Mem I and was incubated for 20 min at RT. Cells were then transfected with this solution in antibiotic-free medium for 24 hours, at final concentration 25 nM siRNA and 0.25% Lipofectamin. After 24 hours, cells were washed with regular medium and let grown until desired day. Cells were taken in pictures using EVOS XL Core for phase contrast, and using Zeiss Axio Observer Z1 for fluorescence.

### Statistical analysis

Statistical tests were conducted in Prism (Version 9.5.1). Shapiro-Wilk test was conducted on experiment results to test their normality. If the normality was confirmed, unpaired two-tailed t-test was used to compare 2 samples, or one-way or two-way ANOVA was used to compare 3 or more samples, followed by either Tukey’s, or Šídák’s multiple comparisons test. If the normality was not confirmed, Mann-Whitney test was used to compare 2 samples, and Kruskall-Wallis was used to compare 3 or more samples, followed by Dunn’s test. All error bars on graph display standard deviation.

## ACKNOWLEDGEMENTS

This work was supported by grants from La Fondation pour la Recherche Medicale (FRM) Amorcage pour les jeunes equipes (AJE20160635985); Fondation ARC pour la Recherche sur le Cancer (PJA20181208104); Agence Nationale de la Recherche (ANR) ANR-19-CE13-0023 and ANR-22-CE14-0062; Ligue Contre le Cancer - Equipe Labellisée 2024, and Institut National du Cancer (INCA-PLBIO23-004-2023-134) (all to W.M. K.). A.K was supported by a fellowship from Eur IMCBio – University of Strasbourg and the Fondation ARC pour la Recherche sur le Cancer, France fourth-year PhD fellowship. D.S.G was supported by the National Research Fund, Luxembourg (FNR) (AFR PhD, Application ID:14584624) and the University of Strasbourg. L.D is funded by an École Doctorale fellowship, University of Strasbourg, France. The work was also supported by an institutional grant of the Interdisciplinary Thematic Institute IMCBio+, as part of the ITI 2021-2028 program of the University of Strasbourg, CNRS and Inserm, was supported by IdEx Unistra (ANR-10-IDEX-0002), and by SFRI-STRAT’US project (ANR-20-SFRI-0012) and EUR IMCBio (ANR-17-EURE-0023) under the framework of the France 2030 Program. The French National Research Agency (ANR) supported the generation, breeding, and phenotyping of mutant mice reported here through the Programme d’Investissement d’Avenir under contract INBS PHENOMIN (ANR-10-INBS-07) grant under the frame programme Investissement d’Avenir ANR-10-IDEX-0002-02. Sequencing was performed by the GenomEast platform, a member of the “France Genomique” consortium (ANR-10-INBS-0009). The funders had no role in study design, data collection and analysis, decision to publish, or preparation of the manuscript.

---

**Extended Data Figure 1.**
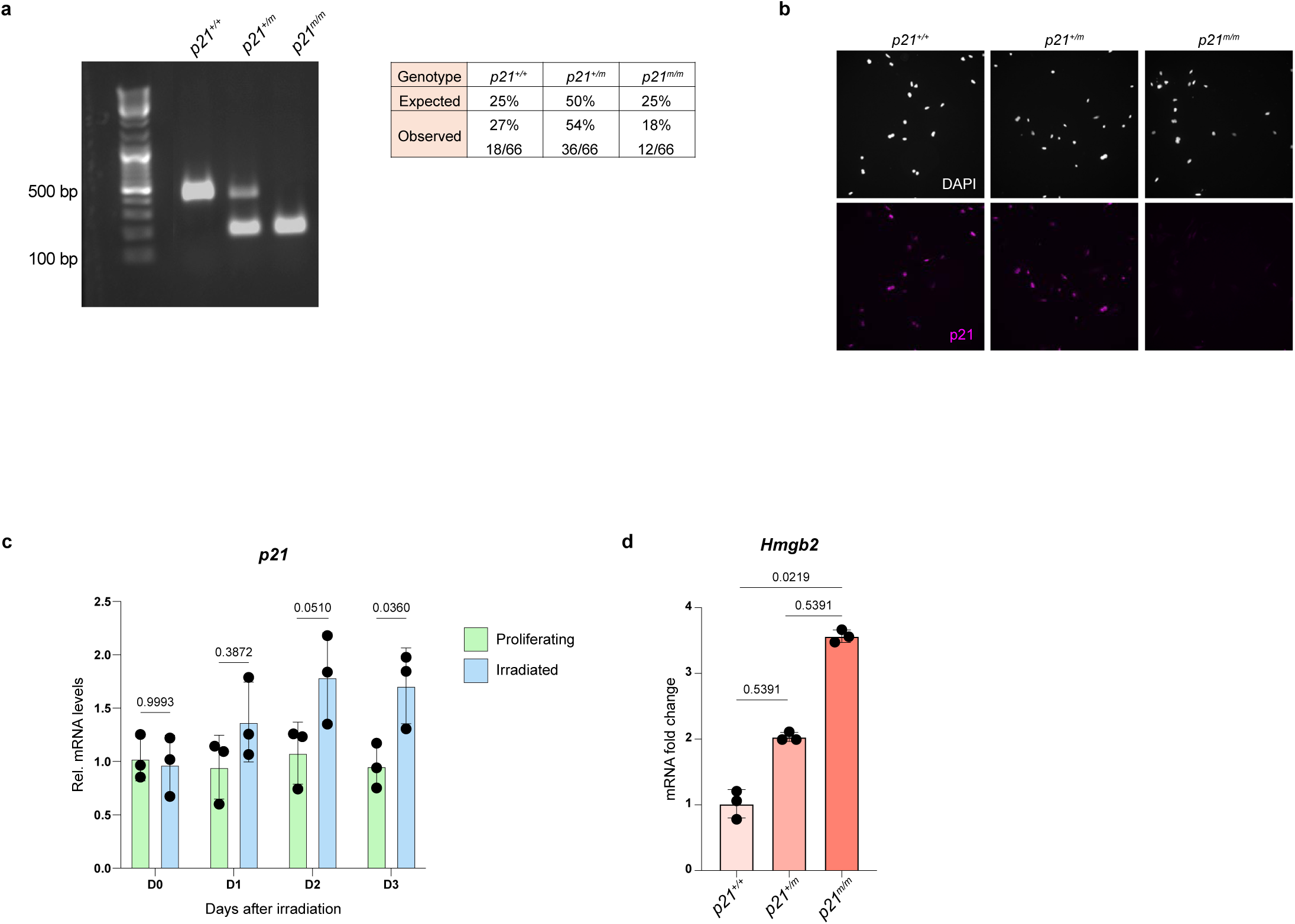
Validation of p21-mCherry-CreERT2 cells. **a**, Left, Target-specific PCR amplification of wild-type, and p21-mCherry-CreERT2 allele in *p21^+/+^*, *p21^+/m^* and *p21^m/m^* mice. Right, Birth ratio analysis of *p21^+/+^*, *p21^+/m^* and *p21^m/m^* mice after intercrosses, and the expected Mendelian ratio. **b**, Immunofluorescence staining for p21 in proliferating *p21^+/+^, p21^+/m^* and *p21^m/m^* MDFs. Scale bars: 200 µm; n=3. **c**, Relative mRNA levels of *p21* levels in D0 to D3 irradiated *p21^+/+^* MDFs, normalized to Rplp0 and control cells before irradiation; n=3, two-way ANOVA test followed by Šídák’s multiple comparisons test. **d**, Relative mRNA levels of *Hmgb2* in *p21^+/+^, p21^+/m^* and *p21^m/m^* MDFs, 2 days after irradiation, normalized to Rplp0 and average of *p21^+/+^*; n=3, Kruskall-Wallis test followed by Dunn’s test.

**Extended Data Figure 2.**
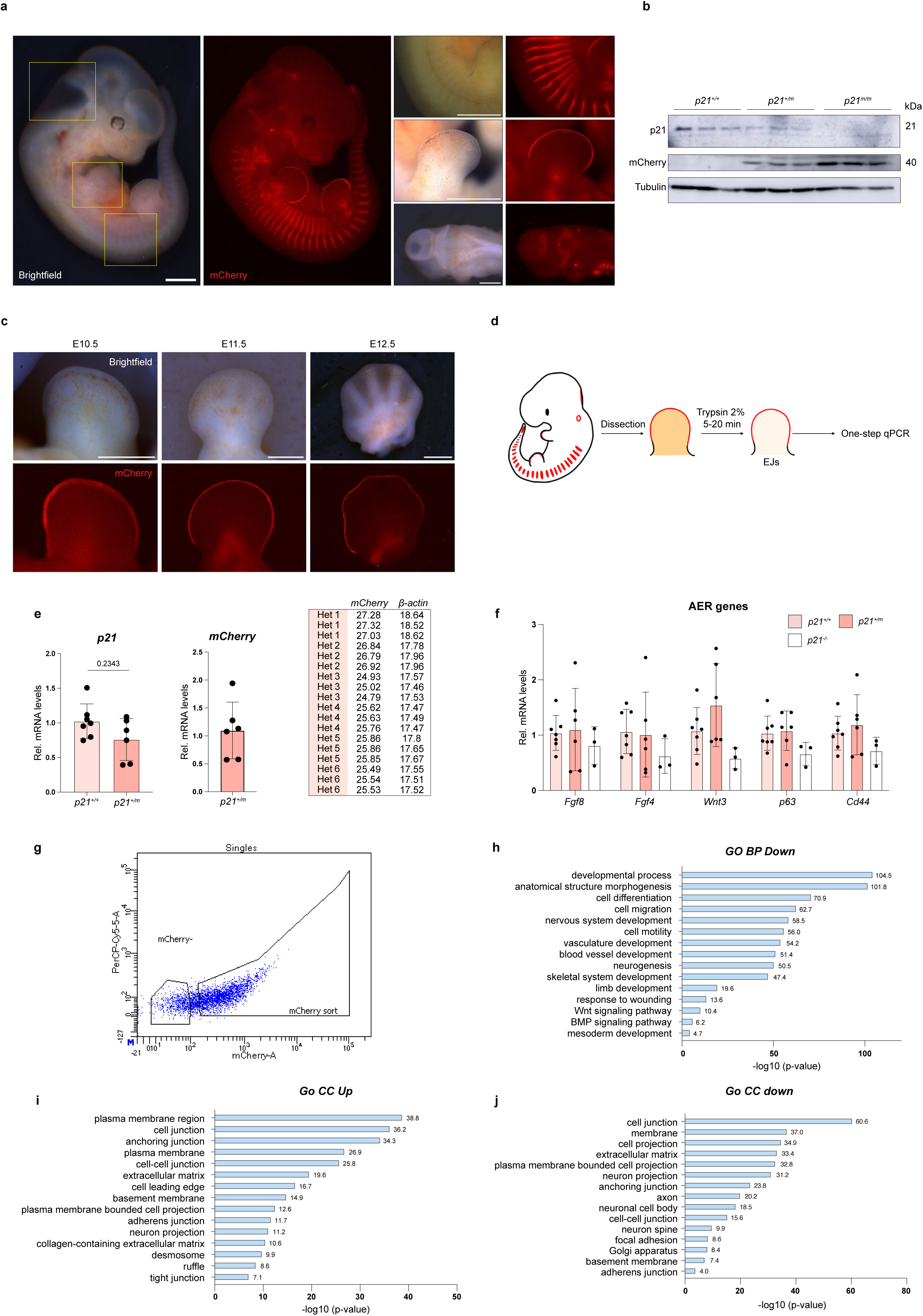
Further analysis of AER and p21-mCherry-CreERT2 mouse model. **a**, Images of live *p21^m/m^* E11.5 embryos. Left pictures showing the pattern of mCherry in whole embryo. Right pictures correspond to high magnification pictures of organs highlighted by yellow boxes; top: somites; middle: forelimb; bottom: hindbrain in dorsal view. n>20, scale bar: 1 mm. **b**, Western Blot for p21, mCherry and Tubulin in whole *p21^+/+^*, *p21^+/m^* and *p21^m/m^*embryos; n=3. **c**, Images of live *p21^+/m^* limbs from E10.5, E11.5 and E12.5 embryos, brightfield (top row) and mCherry fluorescence (bottom row), scale bar: 500 µm. E11.5 mCherry picture was used in figure 2. **d**, Schematic of the dissection protocol to obtain ectodermal jackets (EJs) used for One-step qPCR analysis. **e**, Left, Relative mRNA levels of *p21* in dissected EJs from *p21^+/+^* and *p21^+/m^* E11.5 embryos, normalized to *ß-actin* and compared to *p21^+/+^* samples; n=6-7 from 6-7 different litters; Mann-Whitney test. Middle, Relative mRNA levels of *mCherry* in dissected EJs from *p21^+/m^* E11.5 embryos, normalized to *ß-actin* and average of *p21^+/m^* samples. No detectable mCherry mRNA in *p21^+/+^* embryos. n=6 from 6 different litters. Right, raw CTs detected of *mCherry* and *ß-actin* in *p21^+/m^* EJs samples. **f**, Relative mRNA levels of AER genes, *Fgf8*, *Fgf4*, *Wnt3*, *p63* and *Cd44* in dissected EJs from *p21^+/+^*, *p21^+/m^* and *p21^-/-^* E11.5 embryos. Normalized to ß-actin and compared to *p21^+/+^* samples; two-way ANOVA followed by Tukey’s multiple comparison, no significant difference (p-value > 0.05) detected between conditions for all genes, except between *p21^+/m^* and *p21^-/-^* for *Wnt3* (p-value=0.0174). n= 3-7 from 3-7 different litters. **g**, Fluorescent profile of mCherry in *p21^+/m^* EJs and example of gating strategy for the sorting of mCherry-positive and mCherry-negative cells. **h**, Significant selected GO BP terms of downregulated genes (absolute fold change ≥1.5 and adjusted p-value <0.05) in the AER. **i**, Significant selected GO Cellular Components (CC) terms of upregulated genes (absolute fold change ≥1.5 and adjusted p-value <0.05) in the AER. **j**, Significant selected GO CC terms of downregulated genes (absolute fold change ≥1.5 and adjusted p-value <0.05) in the AER.

**Extended Data Figure 3.**
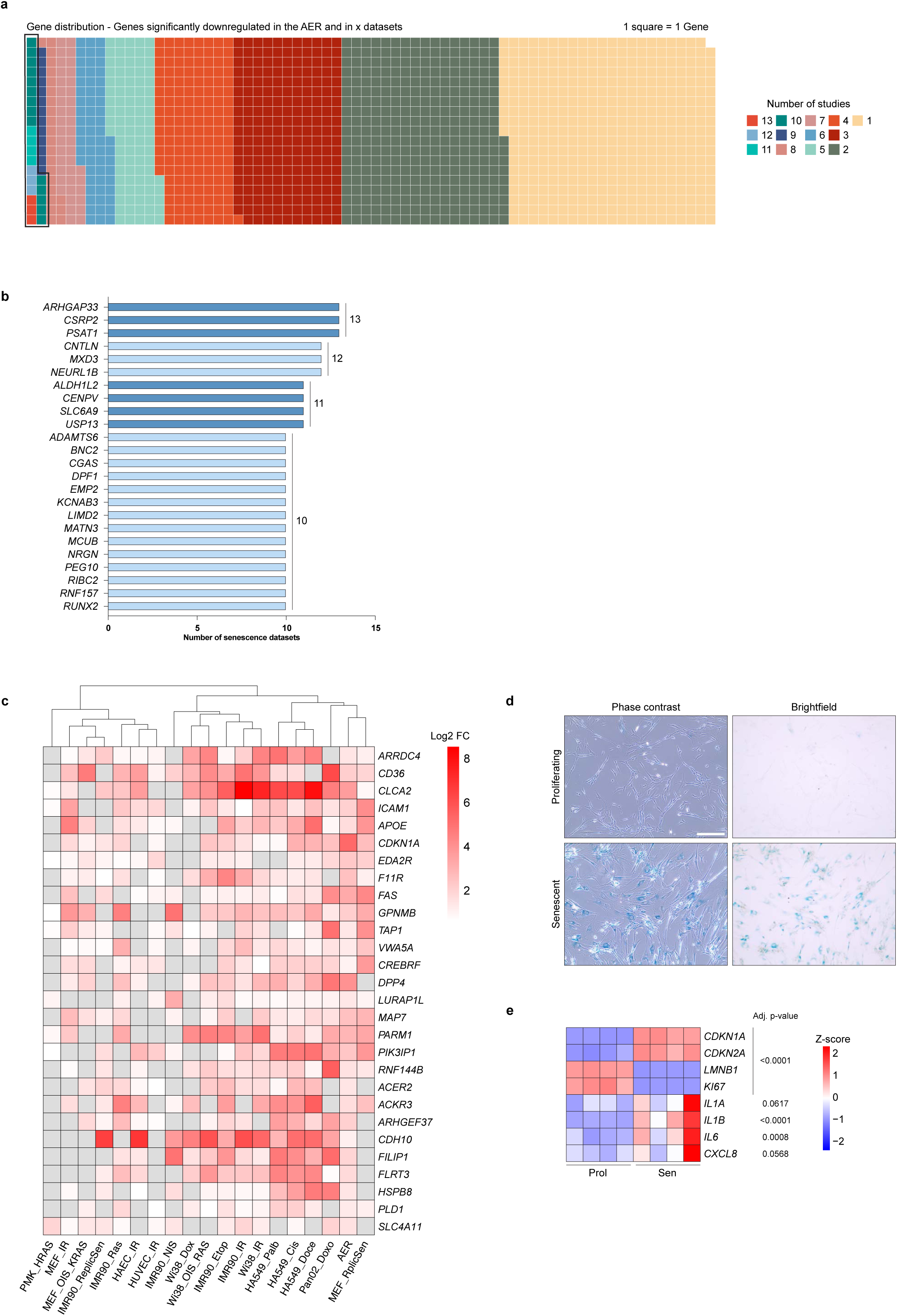
Identification of downregulated genes common to senescence states. **a**, Waffle plot showing the distribution of genes significantly down in the AER and in other *in vitro* senescence studies. 1 square = 1 gene, and each color corresponds to the number of studies in which the gene is downregulated. **b**, Top 24 genes obtained by the meta-analysis and the number of *in vitro* senescence studies in which they are downregulated. **c**, Heatmap of log_2_ FC expression of significantly upregulated top 28 genes in each dataset of the comparative study, row scaling. **d**, SA-ß-Gal staining of proliferating and D10 doxorubicin-induced senescent IMR-90 cells. Scale bar, 300 µm. **e**, Heatmap of normalized reads of classical senescent genes in proliferating and senescent doxorubicin-induced IMR-90 cells, row scaling.

**Extended Data Figure 4.**
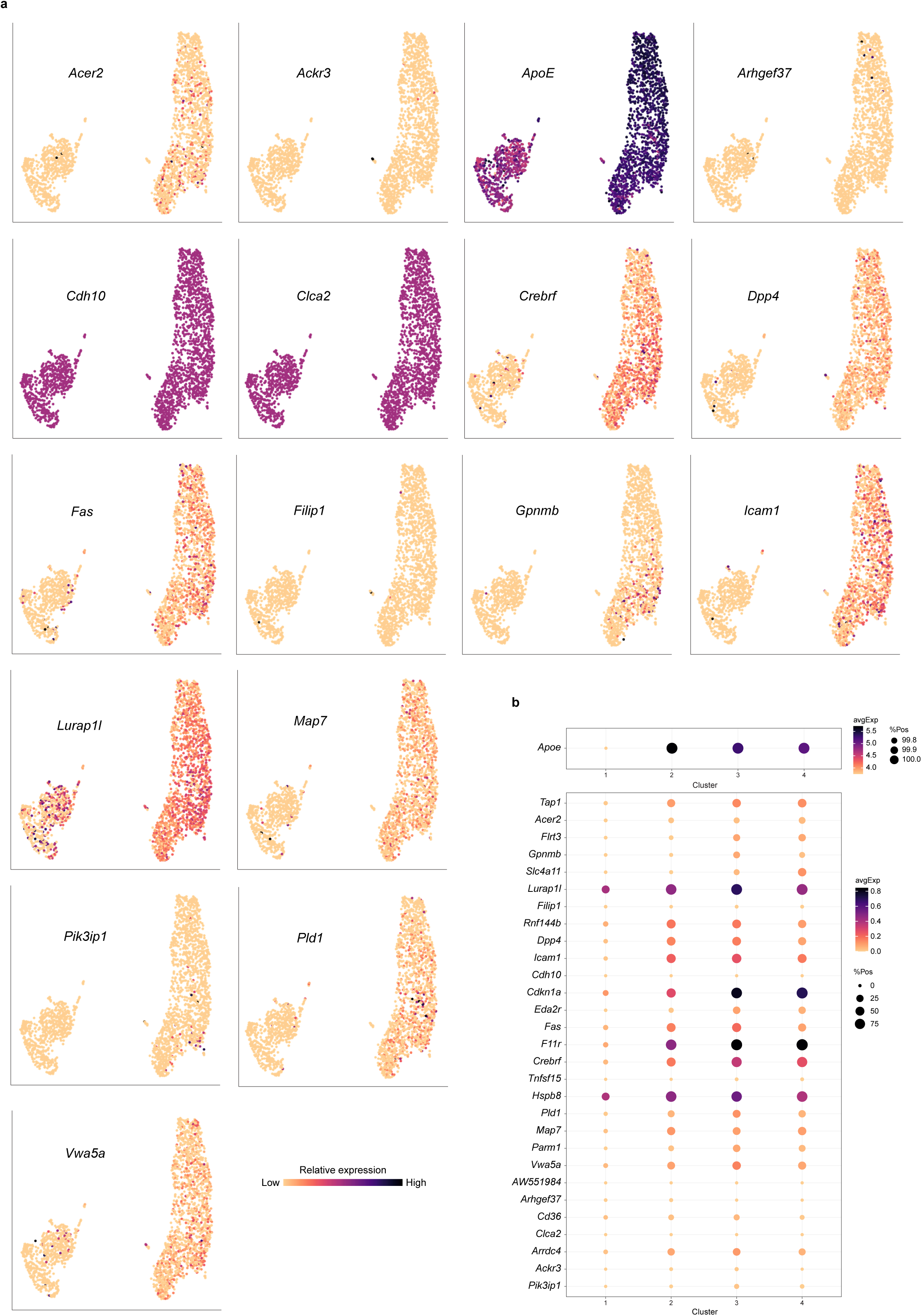
Gene expression of select senescence-associated genes in oncogene-induced senescence in liver cells. **a,** UMAP embeddings of single-cell-sequenced hepatocytes colored by expression of our genes of interest. **b,** Dot plot representing gene expression and percentage of positive cells of our genes of interest in each cluster. Apoe shown in a separate dot plot due to high expression.

**Extended Data Figure 5.**
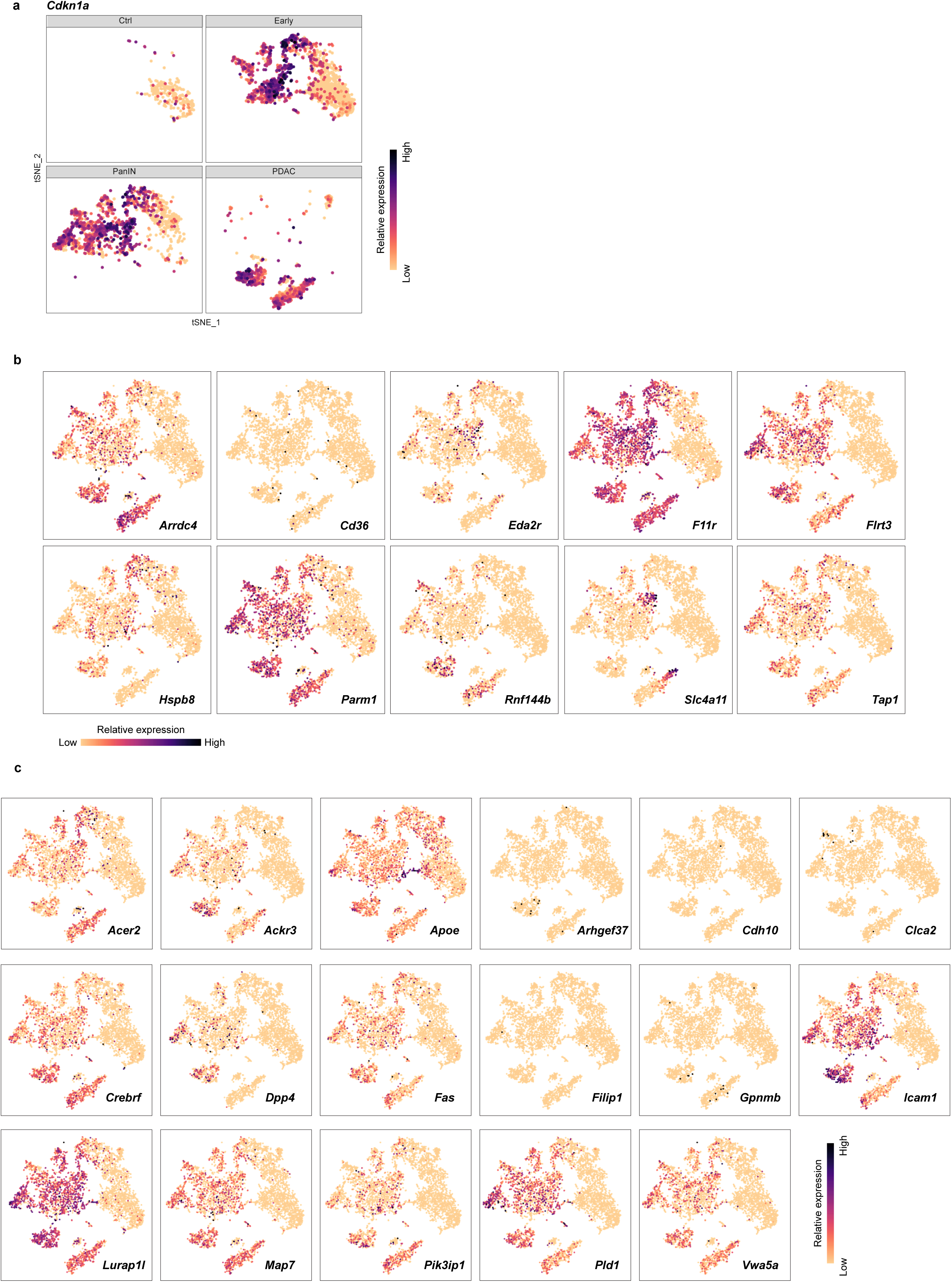
Gene expression of select senescence-associated genes in oncogene-induced senescence in endogenous Kras^G12D^-driven pancreatic tumor model. **a**, *t*-SNE projections colored by *Cdkn1a* expression and separated by sample of origin. **b-c**, *t*-SNE projections colored by expression of (**b**) 10 genes of interest selected for further knock-down, and (**c**) 17 genes of interest upregulated in the AER and in more than 12 senescence datasets.

**Extended Data Figure 6.**
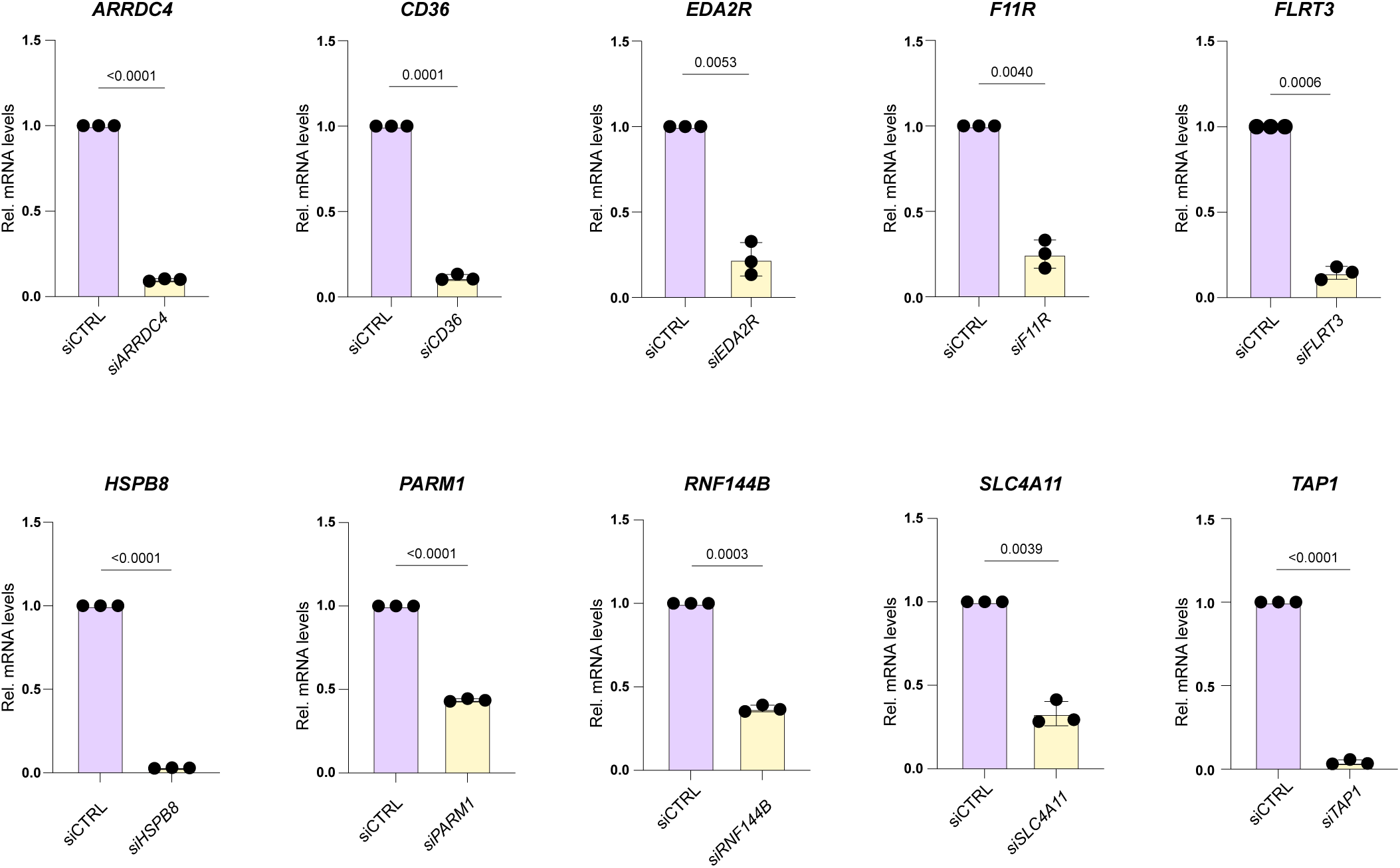
Validation of siRNA knock-down efficiency by RT-qPCR. Relative mRNA levels of each gene of interest targeted by the corresponding siRNA. Normalized to Rplp0 and siCTRL of each experiment. n=3, Welch’s t-tests.

**Extended Data Figure 7.**
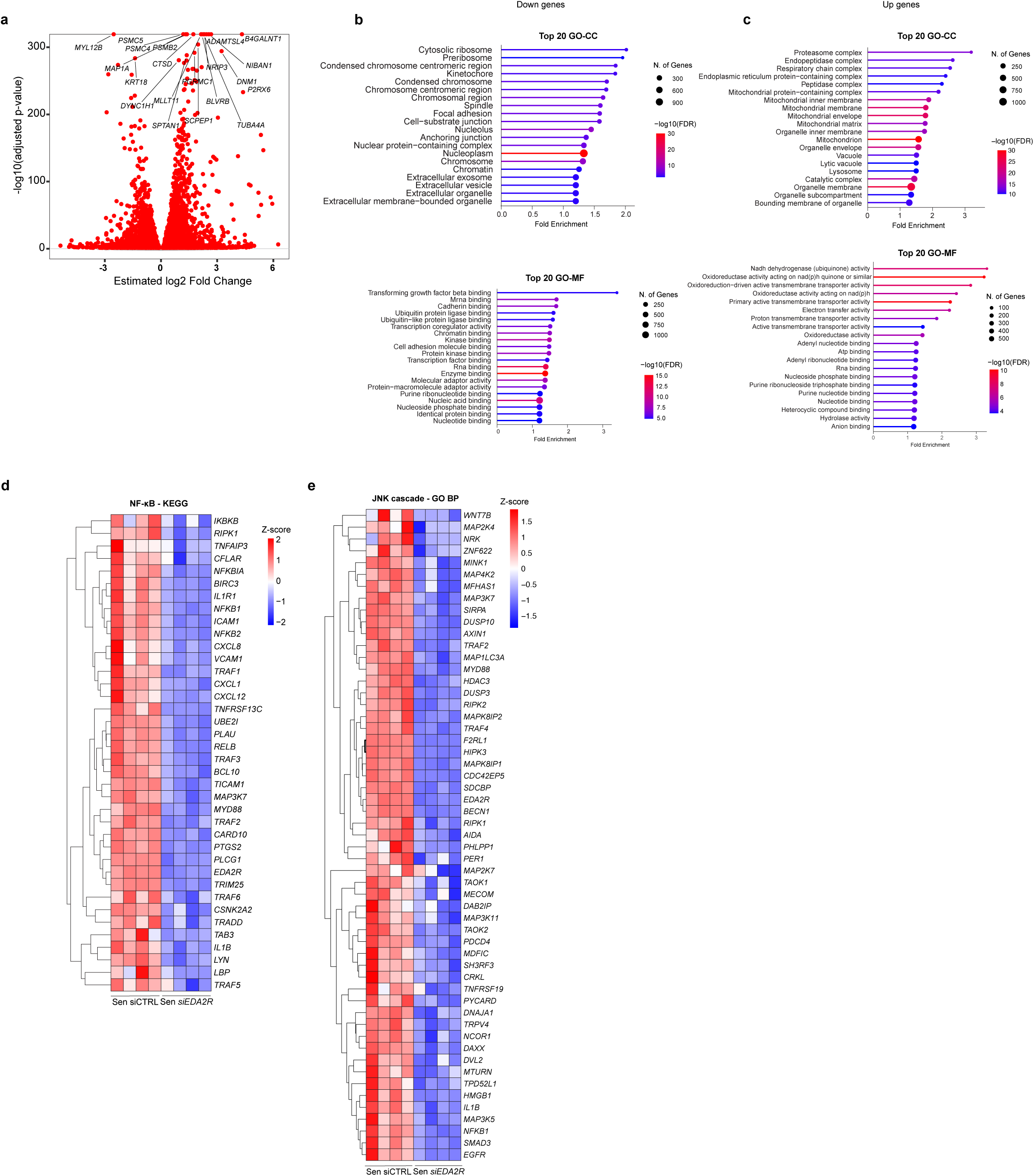
Identification of pathways associated with gene-expression changes upon EDA2R knockdown in senescent cells. **a**, Volcano plots of upregulated and downregulated genes from senescent siCTRL vs siEDA2R and proliferating siCTRL vs siEDA2R comparison (significance determine by adjusted p-value <0.05). **b, c**, Top 20 terms determined by GO CC and GO Molecular Function (MF) pathway analysis of downregulated (**b**) and upregulated (**c**) DEGs common to proliferating and senescent condition (adjusted p-value <0.05) obtained by siCTRL vs siEDA2R comparison. **d**, Heatmap of normalized reads of NF-κB-related genes differentially expressed between senescent siCTRL and senescent siEDA2R samples, from NF-κB KEGG term gene lists. **e**, Heatmap of normalized reads of JNK cascade-related genes differentially expressed between senescent siCTRL and senescent siEDA2R samples, from JNK cascade GO BP term gene lists.

## SUPPLEMENTARY TABLES

**Supplementary Table 1: "**DevSen", all DEGs for which adj. p-value < 0.05

***Supplementary Table 2:*** siEDA2R RNA-seq, all DEGs for which adj. p-value <0.05

***Supplementary Table 3:*** Pathway analysis results for siEDA2R RNA-seq, for the selected terms that are shown in the figures

**Supplementary Table 4:** List of primer sequences

